# Elevated CO_2_ Reshapes the 3D Chromatin Landscape of *Arabidopsis thaliana*

**DOI:** 10.1101/2025.11.20.689512

**Authors:** Scott A. Lewis, Mao Li, Kaushik Panda, Alex Harkess, R. Keith Slotkin, Blake C. Meyers

## Abstract

Atmospheric CO_2_ levels are rising rapidly, yet how plants respond at the genomic level remains poorly understood. The three-dimensional (3D) organization of chromatin in the nucleus is a key regulator of gene expression and environmental response, and changes in chromatin architecture can be transmitted across generations through epigenetic mechanisms. Here, we report that elevated CO_2_ is associated with broad 3D chromatin reorganization in *Arabidopsis thaliana*, spanning scales from chromosome-wide compartment identity to kilobase-resolution Local Chromatin Domains (LCDs) and LCD-Associated Loops (LALs). Using chromatin conformation capture and sequencing (Hi-C) across four mutant backgrounds, we show that these changes exhibit intergenerational persistence in the self-fertilized progeny of plants grown at elevated CO_2_, and that this persistence is sensitive to the Pol V branch of the RNA-directed DNA Methylation (RdDM) pathway rather than to upstream small RNA biogenesis. At kilobase resolution, we annotated 2,279 LCDs whose borders are enriched for transposable element loci occupied by the chromatin-silencing members Pol V and LHP1, and near a third of LCD borders are LALs. Elevated CO_2_ is associated with LAL attenuation and altered transcript abundance at LCD borders marked by H3K27me3, implicating coordinated RdDM and Polycomb group (PcG) activity in the chromatin response to elevated CO_2_. Integrated transcriptome and DNA methylation analysis across all five genetic backgrounds revealed widespread changes in gene and transposable element abundance. These findings support a model in which CO_2_-associated chromatin reorganization confers transcriptional plasticity through persistent chromatin states shaped by the RdDM and PcG pathways.

## Introduction

Atmospheric carbon dioxide (CO_2_) levels have increased rapidly over the last 60 yrs (Keeling et al., 2017), with moderate projections estimating atmospheric levels will exceed 700 ppm within the next 80 yrs (Riahi et al., 2011). Investigating how plants will acclimatize to a high CO_2_ environment is essential. As primary consumers of CO_2_ and the foundation of agricultural output, plants play a critical role in the carbon cycle. While Free Air CO_2_ Enrichment studies have provided fundamental insights into CO_2_-dependent changes in plant physiology (Ainsworth & Long, 2021), the genomic mechanisms driving plants’ acclimatization to elevated CO_2_ remain less understood. Extensive work demonstrates abiotic stress-induced chromatin states in plants that are inherited by non-stressed progeny (Luna et al., 2012; Panda et al., 2023; Rasmann et al., 2012; Slaughter et al., 2012), consistent with an epigenetic basis for these rapid adaptations. Given the sessile nature of most land plants and persistent increase in atmospheric CO_2_, trans-generational epigenetic reprogramming may offer a highly adaptive response mechanism.

Chromatin structure plays a crucial role in cells’ environmental responses (Liu et al., 2017; Sun et al., 2020; Wang et al., 2023), and chromatin modifications support development and stress acclimatization of multicellular organisms (Cavalli & Heard, 2019; Lämke & Bäurle, 2017). How can cells orchestrate these mechanisms with such sophistication? The interplay between DNA methylation, histone modifications, and the 3D structure of chromatin underlie the plasticity of gene regulation (Deng et al., 2014; Domb et al., 2022), providing a molecular blueprint for retaining epigenetic “memories” of environmental conditions across generations via epigenetic reprogramming (Heard & Martienssen, 2014; Lee et al., 2023; Owen et al., 2023). A significant challenge has been in determining the prevalence and functional consequence of 3D chromatin loop domains in plant genomes. Previous work suggests Pol II-mediated transcriptional activity interfaces with the 3D structure of local chromatin to promote gene expression regulation in mouse (Hsieh et al., 2020), yeast (Eser et al., 2017; Tsochatzidou et al., 2017) and *Arabidopsis* (Sun et al., 2024), indicating a role for 3D chromatin topology in transcriptional programming. Because the compact *Arabidopsis thaliana* genome packs regulatory features in dense configuration relative to larger polyploid genomes, resolving local chromatin conformation at the kilobase scale requires deep sequencing to accumulate sufficient contacts per genomic bin (Liu et al., 2016; Wang et al., 2015).

To investigate these critical properties of the 3D genome, we performed a systematic analysis of *Arabidopsis* grown under elevated CO_2_, a condition previously reported to produce a heritable accelerated growth phenotype (Panda et al., 2023). We used chromatin conformation capture (Hi-C) with the Sau3AI enzyme, selected because its recognition site overlaps a cytosine that can be methylated, allowing contact frequency to be related to the local methylation state, here termed methylation-topology. This method was informative for characterizing the direct effect of elevated CO_2_ on DNA methylation and how it relates to 3D genome structure and was highly reproducible (see Methods). Through deep sequencing, we obtained contact maps with an average maximal resolution exceeding 1 kilobase (Kb) and used these maps to model CO_2_-dependent global chromatin architecture rearrangements overlapping epigenetically modified loci. At the kilobase scale, we annotated topological domains of local chromatin loops that are positioned over developmental gene clusters. At a subset of transcriptionally reprogrammed loci, elevated CO_2_ was associated with attenuated or absent chromatin looping coincident with local changes in 5mC, a pattern consistent with chromatin fiber decompaction. Chromosome-wide interaction profiles revealed a distinct chromatin expansion protocol that was reinforced and heritable within the progeny of CO_2_-exposed plants. Together, our findings identify the interaction between 3D structure of chromatin and key chromatin-altering proteins and pathways as a candidate source of epigenetic plasticity in plant acclimatization to elevated CO_2_.

## Results

### Elevated CO_2_ is associated with global chromatin decondensation

The structure and function of the genome is modeled by its 3D conformation. To assess global rearrangements in chromatin architecture, we used Sau3AI-based chromatin conformation capture (Hi-C), in which restriction-site methylation was profiled using matched Enzymatic Methyl-seq (EM-seq) data to account for methylation-dependent cut-site recovery (see Methods) to model the structure of the 3D genome in plants grown at atmospheric vs. 1,000 ppm CO_2_ (Fig. 1A). We also investigated the progeny of plants grown at high CO_2_(“High”), which were returned to ambient CO_2_ levels by seed descent (“pHigh”) (Fig. 1A). Examining the frequency of chromatin interactions between heterochromatic islands at progressively higher resolutions (100 Kb, 20 Kb, 10 Kb, 5 Kb) revealed a molecular phenotype characterized by CO_2_-mediated attenuation of long-range chromatin loops, consistent with chromatin decondensation (Fig. 1B). To profile this effect, we quantified the Interaction Decay Exponent (IDE), which revealed significantly lower genome density across whole chromosomes, pericentromeres, and chromosomal arms (Fig. 1C, Supplementary Fig. S1). Differences in contact-decay between conditions were assessed using a linear model incorporating batch and genomic distances as covariates, with FDR correction applied across pairwise comparisons within each genotype (see Methods); significant comparisons are indicated within the plot and color-coded by condition (Fig. 1C). The interaction decay profiles indicate that the CO_2_-associated change in contact frequency are most pronounced within a subset of pairwise loci separated by less than approximately 1 Mb (Fig. 1C, upper y-axis). To quantify the impact of elevated CO_2_ on global 3D chromatin structure, we simulated 3D models of the genome to profile genome-wide changes in chromatin density (Fig. 1D, Supplementary Fig. S2A). The models recapitulate known features of *Arabidopsis* nuclear organization, such as the extension of chromosome arms from dense chromocenters associated with transcriptionally silent constitutive heterochromatin. The concordant slopes between decay profiles in WT conditions at the macro level are consistent with the broad change in chromatin architecture being primarily a function of chromatin density (Fig. 1B-D). The widespread decondensation of chromatin fibers we infer from Hi-C may increase accessibility to chromatin-interacting proteins in both *cis* and *trans*, a possibility that targeted assays would test directly.

**Figure 1.**
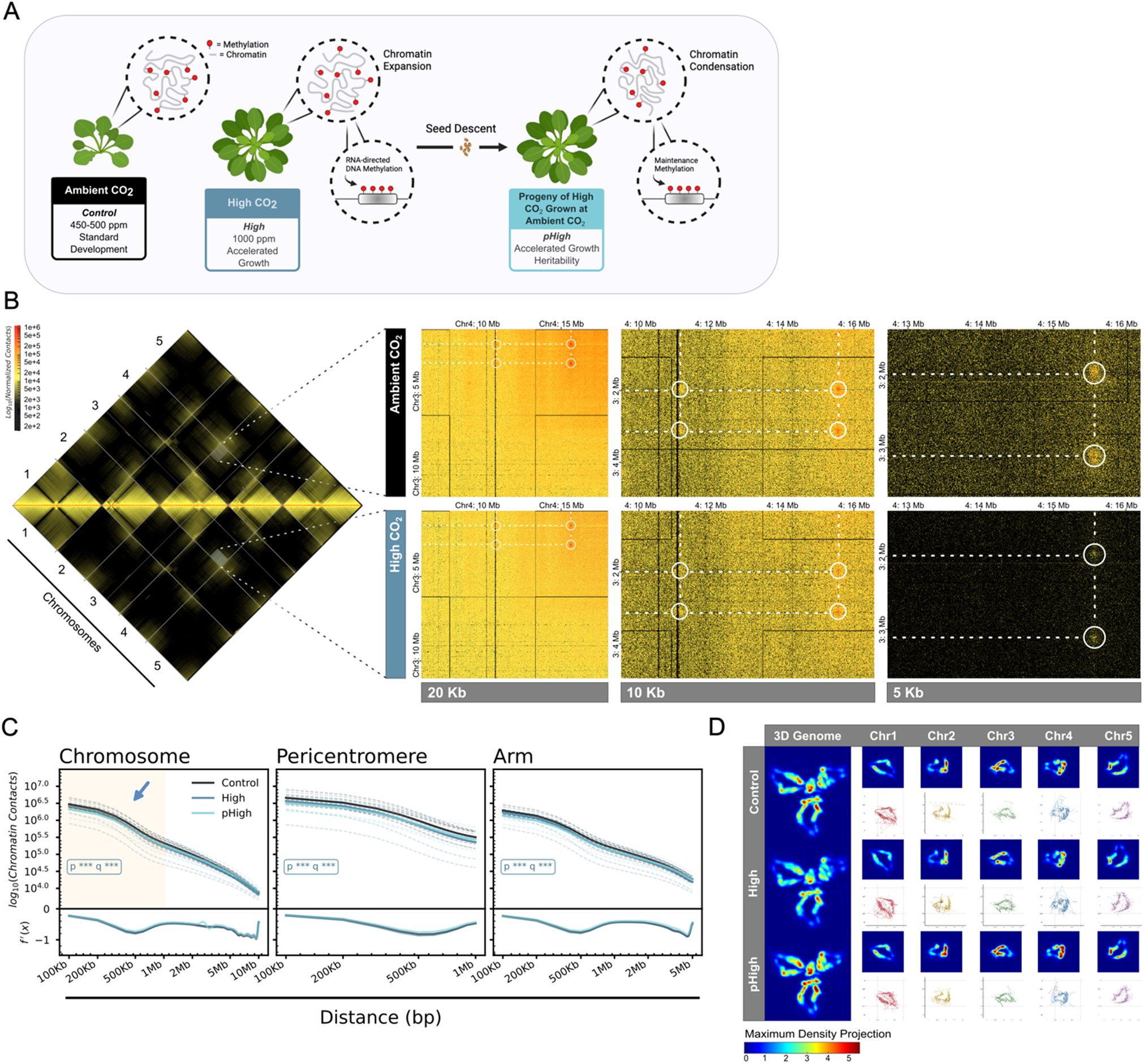
Elevated CO_2_ is associated with global chromatin decondensation and altered 3D genome organization. **(A)** Proposed model of CO_2_-associated chromatin reorganization and experimental design. Wild-type (WT) *Arabidopsis thaliana* plants (Col0 ecotype) were grown at ambient CO_2_ (Control), elevated CO_2_ (High; 1,000 ppm), or as self-fertilized progeny of High CO_2_ plants returned to ambient conditions (pHigh), representing a single generation of seed descent. Based on the data presented in Panels B–D and subsequent figures, we propose a model in which elevated CO_2_ is associated with chromatin expansion and altered DNA methylation and chromatin states at developmentally associated loci in High CO_2_ plants with distinct chromatin expansion patterns in pHigh progeny, consistent with a degree of transgenerational epigenetic memory. **(B)** Split genome-wide view of the *A. thaliana* 3D genome (100 Kb resolution, left). Control (top) and High CO_2_ (bottom) show inter-chromosome contacts. HiGlass browser snapshots (right) for long-range chromatin loops formed between KNOT-Engaged Elements on chromosome 3 and chromosome 4 at 20 Kb, 10 Kb, and 5 Kb resolution in Ambient CO_2_ (top row) and High CO_2_ (bottom row) WT plants. Dashed circles highlight focal points of altered long-range chromatin contact frequency under elevated CO_2_. Color scale reflects distance normalized contact frequency (KR-balanced). **(C)** Log-transformed contact frequency decay curves as a function of genomic distance (bp) for WT Control (black), High (blue), and pHigh (light blue) plants, stratified by genomic context: whole chromosome (left), pericentromeric heterochromatin (center), and chromosome arms (right). Solid liens indicate maximal resolution achieved by combining non-redundant reads across biological replicates. Dashed lines indicate biological replicates shown individually (see Supplementary Fig. S1 for all genotypes). The flattening of decay curves in High CO_2_ plants is consistent with a broad reduction in local chromatin compaction, most pronounced within a subset of pairwise loci separated by less than approximately 1 Mb. **(D)** Maximum density projection heatmaps derived from 3D genome models of all five somatic *Arabidopsis* chromosomes (left column) and individual chromosomes (Chr1–Chr5) in Control, High, and pHigh WT plants. Models were constructed from whole-genome Hi-C contact matrices at 20 Kb resolution using Max3D (Oluwadare et al., 2018) and registered to a common reference via Procrustes alignment. Higher density values (warmer colors) reflect regions of increased chromatin compaction; CO_2_-dependant shifts in density distribution are consistent with global changes in chromatin architecture across conditions.

To investigate mechanisms associated with CO_2_-dependent changes in genome architecture, we analyzed chromatin conformation in genetic backgrounds disrupting specific chromatin-silencing pathways. This set includes CHROMOMETHYLASE 2 and 3 (*cmt2/3*), which maintain non-CpG DNA methylation; DECREASE IN DNA METHYLATION 1 (*ddm1*), an ATPase required for chromatin remodeling; and the specialized RNA polymerases IV and V (*nrpd1* and *nrpe1*), both essential for RNA-directed DNA methylation (RdDM). Pol IV primarily initiates the production of 24-nt small interfering RNAs (siRNAs) that guide DNA methylation, whereas Pol V transcribes scaffold RNAs that recruit methylation machinery to targeted genomic loci, reflecting complementary yet distinct roles in the RdDM pathway. Notably, *nrpe1* is specifically linked to an enhanced heritability of the CO_2_-associated accelerated growth phenotype (Panda et al., 2023)

Two distinct response patterns emerged across genotypes. In *cmt2/3*, *ddm1*, and *nrpe1*, IDE contact frequency increased in High relative to paired controls and decreased in pHigh, indicating a transient increase in genome density in the treated generation followed by expansion in the progeny (Supplementary Fig. S1). In WT and *nrpd1*, contact frequency decreased in both High and pHigh relative to control, indicating expansion in both generations, with the effect further enhanced in *nrpd1* (Supplementary Fig. 1). Principal Component Analysis (PCA) was used to directly compare global differences in chromatin conformation across all groups. PC1 and PC2 explained 38.67% and 21.03% of the total variance, respectively (Supplementary Fig. S2B). Within each genotype, PC1 captured the primary difference in genome-wide conformation between High and pHigh relative to control. Notably, the global conformation profile of *nrpe1* High preferentially clustered with *cmt2/3* Control, consistent with elevated CO_2_ disrupting the maintenance of *de novo* Pol V RdDM by CMT2/3. However, *cmt2/3* High shows condensation, instead of the expansion observed in WT High, indicating that additional pathways involved in elevated CO_2_-mediated dysregulation remain to be investigated.

Chromatin decondensation appears to involve interplay between distinct pathways. The observation that *nrpd1* conserves the WT response, with expansion in both generations, is consistent with elevated CO_2_-dependent Pol IV attenuation, genetic ablation of Pol IV adds little to a conformational phenotype that CO_2_ treatment alone produces. By contrast, loss of Pol V (*nrpe1*), maintenance methylation (*cmt2/3*), or chromatin remodeling (*ddm1*) redirects to a biphasic response, producing condensation in the treated generation before expansion in the progeny. The observation that *nrpe1* pHigh progeny show both the strongest chromatin expansion and enhanced heritability of the accelerated growth phenotype (Panda et al., 2023) links the magnitude of the conformational change in the progeny generation to the transmitted phenotype. The correlation between WT and *cmt2/3* conformations at elevated CO_2_ is consistent with a model in which chromatin remodeling by DDM1 contributes to the deposition of Pol V-mediated *de novo* DNA methylation signatures and their heritable maintenance by CMT2/3 in dense heterochromatin, facilitating chromatin decondensation.

### Chromatin decondensation is accompanied by a reorganized compartmentalization program

The nuclear territories of chromosomes are spatially resolved into compartments of transcriptionally active chromatin (‘A compartments’) and transcriptionally repressed chromatin (‘B compartments’) (Domb et al., 2022; Lieberman-Aiden et al., 2009). To examine how elevated CO_2_ impacts chromatin structure and genome activity, we analyzed genome-wide chromatin A/B compartments at 20 Kb resolution. This analysis identified broad changes in genome spatial organization (Fig. 2A-B). Compared to the Control group, WT High and pHigh plants showed substantial changes in chromatin structure: inactive heterochromatin near the centromeres and in the pericentromeres interacted more frequently with chromosome arms which are composed primarily of euchromatin, consistent with a reorganization of compartment domains under elevated CO_2_. Increased heterochromatin–euchromatin interactions were maintained in the progeny generation (pHigh), consistent with heritable transitions in chromatin state (Fig. 2B).

**Figure 2.**
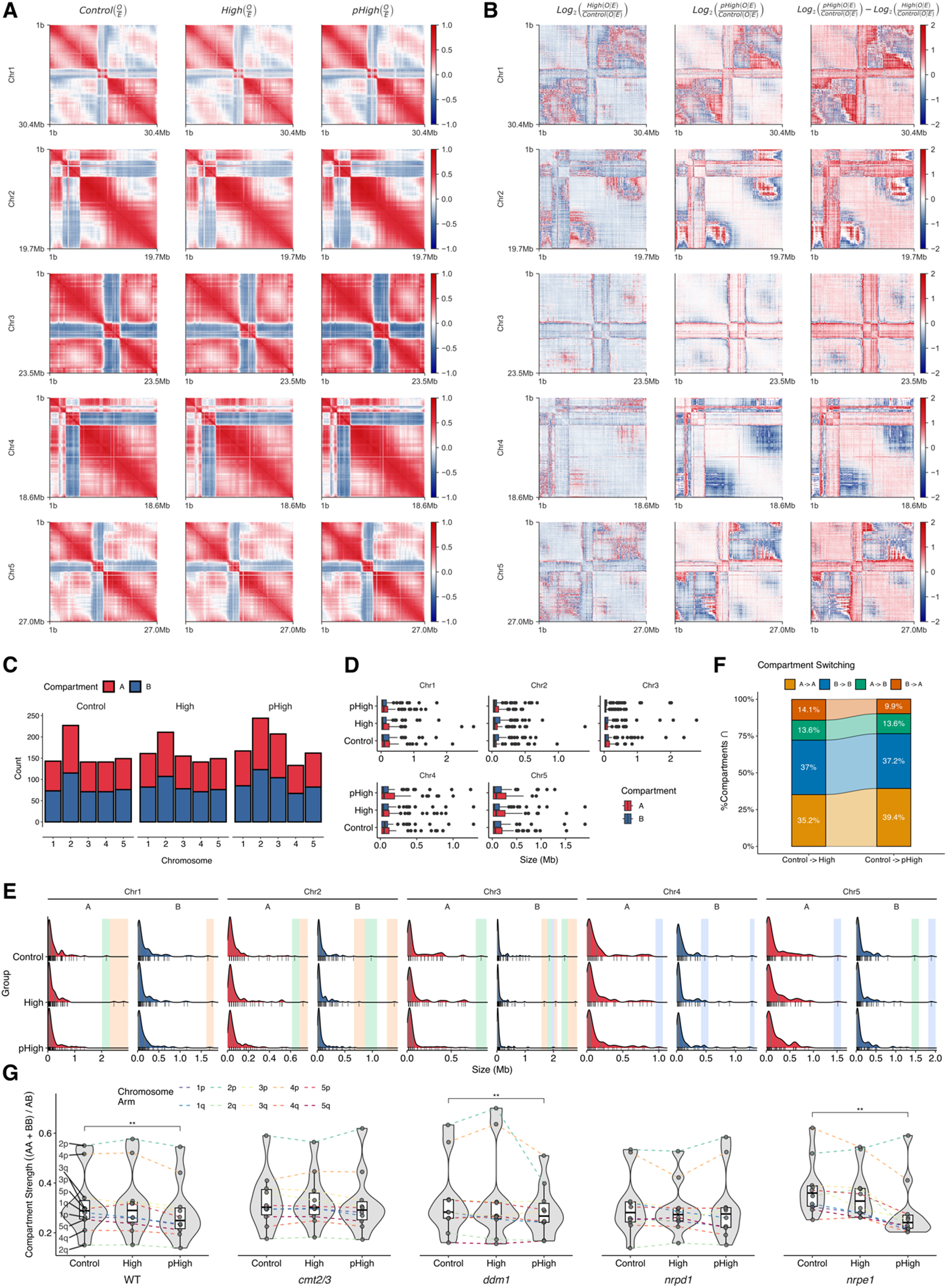
Elevated CO_2_ is associated with broad changes in chromatin compartment organization and genome activity. **(A)** Genome-wide correlation matrices of chromatin interactions binned at 20 Kb resolution, shown per chromosome for wild-type (WT) plants grown at ambient CO_2_ (Control), elevated CO_2_ (High; 1,000 ppm), and progeny of High CO_2_ plants returned to ambient conditions (pHigh). Positive correlations are indicated in red and negative correlations in blue. **(B)** Log_2_ fold-change in chromatin interaction correlation per chromosome for High and pHigh plants relative to paired internal Control plants, highlighting regions of gained (red) and lost (blue) intra-chromosomal interaction frequency. **(C)** Quantification of annotated A (transcriptionally active; red) and B (transcriptionally inactive; blue) compartments per chromosome in each WT condition. Compartments were annotated using the signed PC3 eigenvector at 20 Kb resolution (see Methods; Supplementary Fig. S3). **(D)** Length distribution of annotated A and B compartments per chromosome in each WT condition. **(E)** Density profiles of compartment length distributions illustrating examples of delayed (blue shading), transient (orange shading), and persistent (green shading) compartment switching. Delayed transitions emerge in pHigh but not High; transient transitions are present in High but revert in pHigh; persistent transitions are maintained across both High and pHigh conditions. **(F)** Alluvial summary of compartment switching between WT Control, High, and pHigh plants, showing the proportion of A and B compartments that switch identity or are maintained across conditions. **(G)** Compartment strength quantified per chromosome arm across all five somatic *A. thaliana* chromosomes in all five genetic backgrounds (WT, *cmt2/3*, *ddm1*, *nrpd1*, *nrpe1*) and CO_2_ treatment groups. Compartment strength is calculated as the ratio of intra-compartment (AA, BB) to inter-compartment (AB, BA) interaction frequency. The x-axis hierarchy denotes genetic background followed by treatment group. Brackets indicate statistically significant pairwise differences in compartment strength (Wilcoxon rank-sum test, P < 0.05, FDR < 0.05. Attenuation of compartment strength in pHigh, *nrpe1*, and *ddm1* is consistent with disruption of RdDM and chromatin remodeling pathways in maintaining chromatin domain integrity.

We classified A and B compartments using PCA, extracting the weighted dimensional loadings oriented by putative open chromatin signals (see Methods). Like the compact *Drosophila melanogaster* genome, the leading eigenvector (PC1) primarily reflected the separation of chromosome arms from centromeric heterochromatin, rather than *bona fide* compartments (Kalluchi et al., 2023; Lucchesi, 2019; Rowley et al., 2017; Sexton et al., 2012); we therefore used the PC3 eigenvector to annotate A and B compartments, because it showed the greatest correspondence with euchromatic and heterochromatic chromatin marks among the leading eigenvectors, and strong per-chromosome Pearson correlation with previously published *Arabidopsis* compartment annotations, whereas PC1 and PC2 were dominated by the arm-centromere axis of each chromosome (Supplementary Fig. S3, see Methods).

In total, 802 distinct compartments were annotated in WT Control plants, spanning less than 1 Mb of linear genomic sequence on average (Fig. 2, C and D). Comparing the WT Control compartments to those of the High and pHigh groups, we identified prominent shifts in compartment size and representation (Fig. 2, C and D). Using the progeny (pHigh) as the heritability readout, we classified compartmental shifts as persistent (present in High and retained in the pHigh progeny; green), transient (present in High but reverted in pHigh; orange), and delayed (absent in High but emerging in pHigh; blue) (Fig. 2E). In WT, 13.6% of A compartments heritably transitioned to B compartments in both High and pHigh (Fig. 2F), while 14.1% and 9.9% of B compartments transitioned to A compartments in High and pHigh, respectively (Fig. 2F).

Compartment strength decreased progressively from Control to High to pHigh in WT, *ddm1,* and *nrpe1*, while *cmt2/3* and *nrpd1* did not show comparable significant attenuation (Fig. 2G). The compartment analyses are consistent with extensive chromatin reorganization under elevated CO_2_ and support broad changes in genome activity. These results are consistent with Pol IV ablation alone (*nrpd1*) not being sufficient in recapitulating the compartment dysregulation identified in WT, *ddm1*, and *nrpe1*, and points to an ensembled mechanism, where at elevated CO_2_ the loss of Pol V (*nrpe1*) produces compartment attenuation independent from, and equally significant with the attenuation observed with the loss of chromatin remodeling complex DDM1 (*ddm1*), as well as within an unperturbed genetic background (WT). Isolated Pol V genetic ablation (*nrpe1*) does not suppress, and may enhance, the accelerated growth phenotype and magnitude of compartment attenuation. Genetic perturbations clarified the general contributions of chromatin remodeling (DDM1) and RdDM (Pol IV–V) to genome compartmentalization, however specific transcription factor binding events and histone modifications were not assayed directly under elevated CO_2_. Their potential contributions to the observed reorganization are addressed below through enrichment analyses using external datasets and should be regarded as candidate associations rather than demonstrated mechanisms.

### Local chromatin domains are characterized by chromatin state heterogeneity

Chromatin structure is hierarchical, with local interaction clusters nested within compartments and the boundaries between them. Given the possibility that broad shifts in chromatin density and compartmentalization could be supported by changes in smaller, locus-specific 3D structures, we sought to characterize LCDs via chromatin state association and regulatory propensity. We used the maximal resolution of our Hi-C libraries to annotate and characterize LCDs and their associated chromatin states across the *Arabidopsis* genome. Using a local minima approach (see Methods), we annotated 2,279 Local Chromatin Domains (LCDs) genome-wide at 1 Kb resolution, under normal conditions. An illustration depicts a working model of how localized chromatin topology interacts with CRE activity near embedded TEs and developmentally associated gene clusters (Fig. 3A). A representative browser view of an LCD over the *FLOWERING LOCUS C* (*FLC*) locus illustrates characteristic features of LCD organization, such as elevated intra-domain contact frequency, flanking regions of reduced contact at domain boundaries, and the presence of TE loci near the borders of these regions (Fig. 3B).

**Figure 3.**
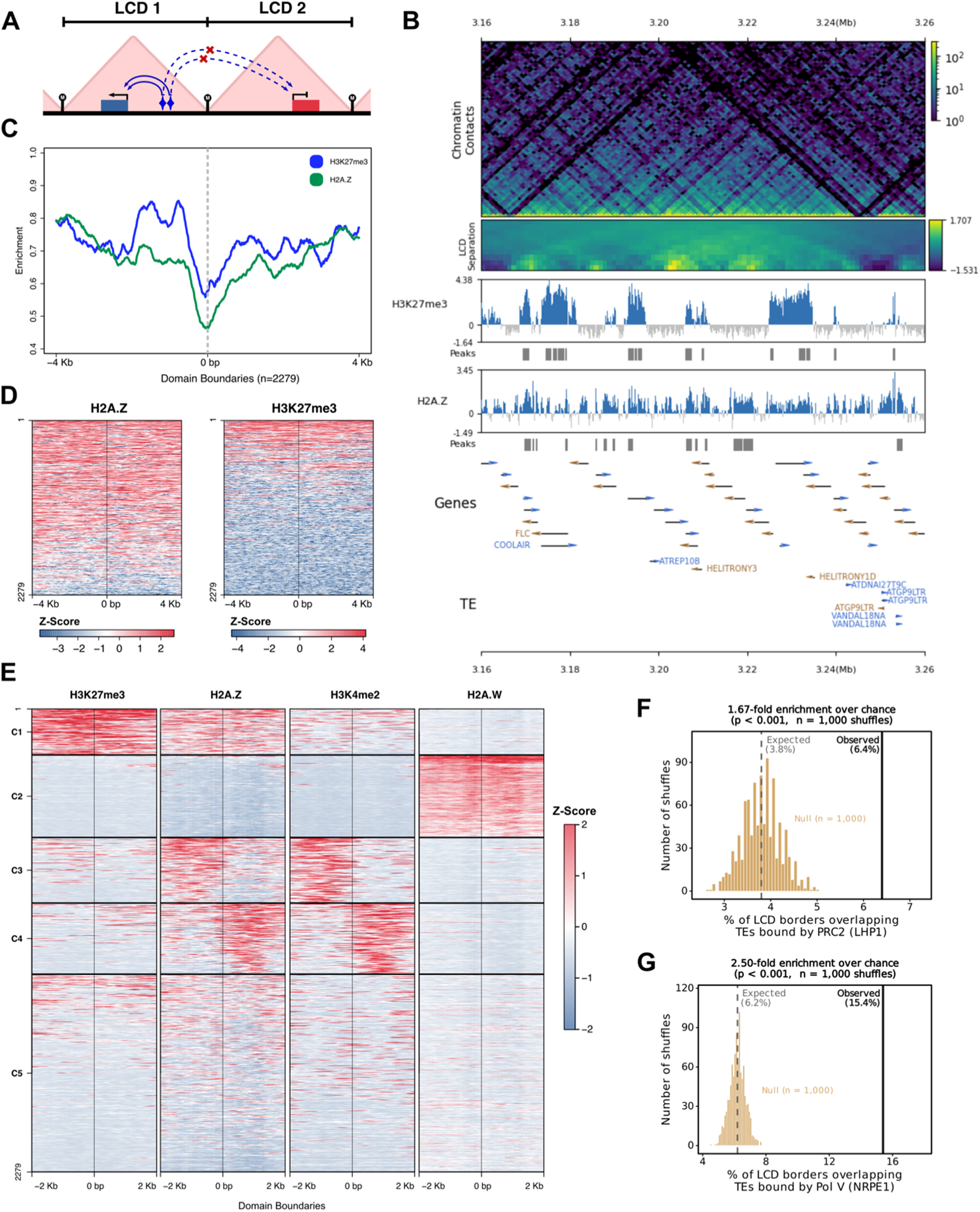
Local Chromatin Domains (LCDs) are associated with characteristic histone mark enrichment and are significantly enriched at borders for transposable element loci bound by the DNA methylation and Polycomb group machineries. (A) Schematic model of an LCD border illustrating the proposed structural organization: silenced loci bound by RdDM and PcG machinery at domain borders anchor chromatin loops over clusters of developmentally regulated genes in the domain interior. CREs harbored within border TEs may regulate flanking gene expression through proximity looping. **(B)** Representative genome browser view of an LCD over the *FLOWERING LOCUS C* (*FLC*) and anti-sense COOLAIR loci on chromosome 5 (Chr5: 3.16–3.26 Mb). Top: KR-normalized Hi-C contact frequency heatmap at 1 Kb resolution (WT Control). LCD boundaries visible as regions of reduced contact frequency flanking the domain. Medium-intensity interactions from the left border region (3.17 - 3.18 Mb) show visible chromatin loops not subject to Brownian motion-induced artefacts after distance normalization (KR-balancing). Middle: LCD separation score track used for domain boundary annotation; negative separation score values delineate local neighborhoods of chromatin interactions. Statistically significant boundaries were annotated as LCD borders (FDR < 0.05). H3K27me3 and H2A.Z ChIP-seq signal (normalized to 1X coverage) with IDR peaks indicated as grey bars below associated distributions. Bottom: gene annotation track showing *FLC*, *COOLAIR*, and neighboring gene loci (not labelled for visual clarity) (orange = positive strand, blue = negative strand). TE loci (ATREP10B, HELITRONY3, HELITRONY1D, ATGP9LTR, VANDAL18NA) are visible in proximity to the LCD borders, consistent with a silent TE-anchored domain boundary structure. **(C)** Metaplot of H3K27me3 (blue) and H2A.Z (green) ChIP-seq enrichment over an 8 Kb window (±4 Kb) centered on all 2,279 LCD borders. y-axis: Z-score, number of standard deviations from the length-weighted mean enrichment of the 1X normalized ChIP-seq signal. H3K27me3 and H2A.Z show enrichment near LCD borders, with H3K27me3 showing a moderate positive correlation with H2A.Z across borders (Pearson r = 0.44; see panel E and Fig. 3D). **(D)** Z-score co-ordered heatmaps of H2A.Z (left) and H3K27me3 (right) ChIP-seq signal over an 8 Kb window centered on all 2,279 LCD borders, ordered by H2A.Z signal. Co-enrichment of both marks is evident at a subset of LCD borders, with 32.6% of borders (743 / 2,279) simultaneously overlapping high-confidence H3K27me3 and H2A.Z IDR peaks as a conservative estimate. **(E)** Z-score co-ordered heatmaps of H3K27me3, H2A.Z, H3K4me2, and H2A.W 1X normalized length-weighted mean ChIP-seq signal enrichment at LCD borders, stratified by five clusters (C1–C5) identified by k-means clustering (k = 5, determined by within-cluster sum-of-squares minimization). Clusters reflect distinct chromatin states at LCD borders: C1–C2 are co-enriched for H3K27me3 and H2A.Z consistent with facultative heterochromatin adjacent to active transcriptional regions; C3–C4 show H3K4me2 enrichment consistent with active euchromatin; C5 is enriched for H2A.W consistent with placement in constitutive heterochromatin (Yelagandula et al., 2014). **(F)** Permutation enrichment test showing that LCD borders are enriched for TE loci occupied by the PcG maintenance factor LHP1 (observed 6.4% vs. 3.8% expected by chance; 1.67-fold enrichment; permutation test, n = 1,000 shuffles, p < 0.001). Amber bars show the null distribution of overlap fractions from 1,000 random shuffles of LCD border coordinates; dashed line indicates the mean expected overlap; solid vertical line indicates the observed overlap. **(G)** Permutation enrichment test showing that LCD borders are enriched for TE loci bound by Pol V (NRPE1) as measured by ChIP-seq (Zhong et al., 2012) (observed 15.4% vs. 6.2% expected by chance; 2.50-fold enrichment; permutation test, n = 1,000 shuffles, p < 0.001). Together, panels F and G are consistent with a model in which RdDM-silenced and PcG-maintained TE loci are structural components of LCD borders, and that CREs within these TEs may regulate flanking gene expression through proximity looping.

To characterize chromatin states associated with LCD borders more broadly, we identified patterns of co-enrichment between key histone modifications and variants using published ChIP-seq datasets (Fu et al., 2022; Liu et al., 2018) (Supplementary Data S3). LCD borders overlapping constitutive heterochromatin were enriched for H2A.W, a histone variant that defines plant heterochromatic regions (Yelagandula et al., 2014), whereas borders outside constitutive heterochromatin showed co-enrichment for tri-methylated histone H3 lysine 27 tri-methylation (H3K27me3), histone variant H2A.Z, and histone H3 lysine 4 di-methylation (H3K4me2). K-means clustering of border mark enrichment profiles (k = 5, clusters C1–C5) revealed distinct chromatin state classes at LCD borders, ranging from H3K27me3/H2A.Z co-enriched (C1) to H3K4me2-enriched (C3–C4) to H2A.W-enriched (C2) (Fig. 3E). 32.6% of LCD borders (742 / 2,279) simultaneously overlapped high-confidence H3K27me3 and H2A.Z IDR peaks within a ±4 Kb window, consistent with co-enrichment of both marks at a substantial subset of euchromatin-associated LCD borders (Fig. 3C-E, Supplementary Figure S4; Pearson r = 0.44). These patterns of histone-mark co-enrichment were not recovered in a random distribution of positionally-shifted LCD borders (Supplementary Fig. S4), consistent with a modular organization in which LCD borders demarcate chromatin environments in both transcriptionally active euchromatin and repressive heterochromatin.

Consistent with a role for RdDM-mediated TE silencing in organizing chromatin domain boundaries, 15.4% of LCD borders (352 / 2,279) overlapped TE loci bound by Pol V (NRPE1) in published ChIP-seq datasets, representing a 2.50-fold enrichment over the 6.2% expected by chance (permutation test, n = 1,000 shuffles, p < 0.001; Fig. 3G). LCD borders were also enriched for loci occupied by the PcG maintenance factor LHP1 (6.4% observed vs. 3.8% expected; 1.67-fold enrichment; permutation test, n = 1,000 shuffles, p < 0.001; Fig. 3F). These border populations were not mutually exclusive. A common set of 50 LCD borders carried both Pol V- and LHP1-bound TE loci, a significant co-occurrence (Fisher’s exact test, OR = 3.16, 95% CI 2.15–4.59, p = 4.5 × 10⁻⁹) indicating that the RdDM and PcG pathways, though mechanistically distinct, converge on a shared subset of LCD borders that shape high-resolution 3D chromatin structure. The joint enrichment of distinct silencing machineries across borders is consistent with a model in which RdDM-silenced and PcG-maintained TE loci contribute to the formation and maintenance of LCD borders. Chromatin-state and TE-silencing signatures at LCD borders, together with the observed domain organization over developmentally associated gene bodies, such as the FLC locus (Fig. 3B), inform a model in which LCD borders delimit local chromatin domains, calibrating the 3D spatial proximity of chromatin fibers near developmentally associated loci (Fig 3A).

### Elevated CO_2_ is associated with changes in local chromatin loop structures

In WT, 32.8% of LCD borders (749/2,279) coincided with chromatin loop anchors identified in WT Control at 1 Kb resolution within ±4 Kb (see Methods), indicating that a substantial number of LCD borders concomitantly serve as structural loop anchors under normal growth conditions, connecting regions upstream of the LCD border to regions downstream of the LCD. We refer to these as LCD-Associated Loops (LALs) and analyzed them as aggregate pile-ups of the contact signal centered on LCDs (Fig. 4A; see Methods). Because LALs are oriented relative to the LCD start, we standardized to orientation of LCD coordinates to a common 5ʹ to 3ʹ convention before aggregation, accounting for the asymmetry of the aggregate signal (see Methods). Of the 749 LALs we identified, 70.9% (531/749) overlapped H3K27me3 IDR peaks within the same window, indicating that the majority of LALs are embedded within H3K27me3-enriched chromatin. LALs were detectable in WT under normal conditions and were reduced in normalized aggregate signal under elevated CO_2_ treatment (Fig. 4A, red dots mark the loop-anchor positions in the pile-up). The mean observed/expected contact frequency at LAL anchor positions in the aggregate pile-up was reduced by approximately 50% in WT High relative to WT Control. Per-replicate aggregate scores and per-locus contact ratios across 897 unique LAL contact loops confirmed this attenuation, 59.8% of individual LALs showed reduced contact frequency in WT High and 63.3% in WT pHigh relative to WT Control, consistent with coordinated loop attenuation across the LAL population rather than concentrated attenuation at a small subset of loci (Supplementary Fig. S5C).

**Figure 4.**
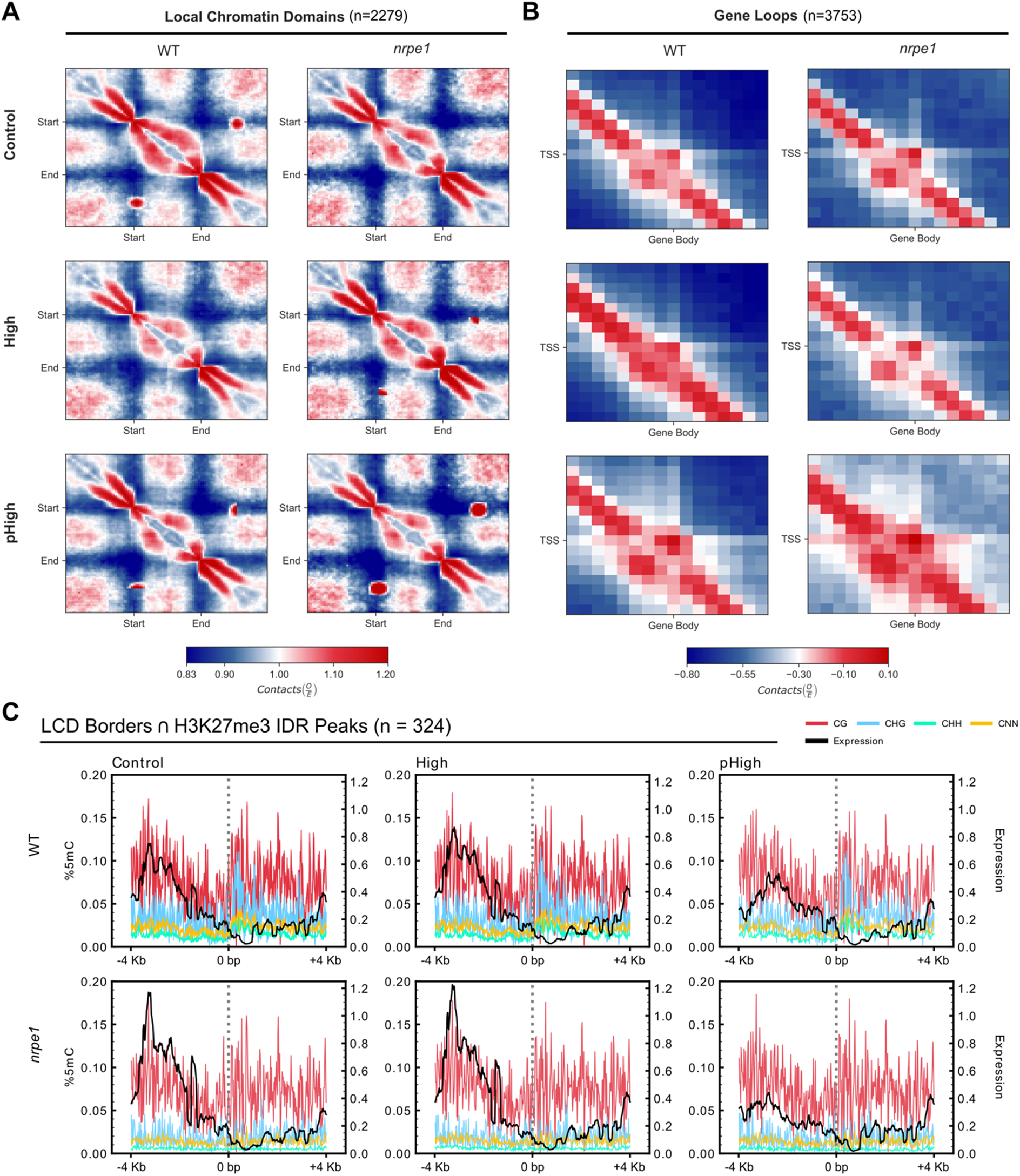
Elevated CO_2_ is associated with changes in local chromatin loop structure at LCD borders and gene loci, particularly in contexts marked by H3K27me3. (A) Aggregate contact frequency maps (pile-ups) over 2,279 annotated LCD regions in WT and *nrpe1* plants under Control, High CO_2_, and pHigh conditions. Contact matrices were generated at 1 Kb resolution and normalized by the mean contact frequency at equivalent genomic distances (see Methods). The characteristic increased contact frequency along the Start–End diagonal and between Start and End anchors reflects the internal organization of LCDs. The red dot visible at the downstream anchor position represents the concentrated contact signal from the 749 LCD borders that coincide with WT Control chromatin loop anchors (LALs; 32.8% of all LCD borders), which accumulates at a consistent position in the oriented pile-up against the broader LCD contact background. In WT High plants, LCD contact frequency is reduced relative to Control, consistent with a CO_2_-associated attenuation of local chromatin loop structure within LCDs. This pattern is partially restored in pHigh, consistent with a degree of transgenerational memory of CO_2_-associated chromatin reorganization. The *nrpe1* mutant shows altered LCD contact patterns under all conditions relative to WT, consistent with a role for Pol V-mediated RdDM in maintaining local chromatin loop structure at LCD border loci. Color scale indicates normalized (KR-balanced) contact frequency (0.55–1.25). **(B)** Aggregate contact frequency maps over 3,753 annotated gene self-loops, defined as chromatin loops where both anchors map to the same gene locus, shown for WT and *nrpe1* plants under Control, High CO_2_, and pHigh conditions. Gene self-loop contact frequency is reduced in WT High and further attenuated in pHigh relative to Control, consistent with CO_2_-associated changes in the organization of local chromatin supporting highly transcribed genes. The *nrpe1* mutant shows increased variability in gene self-loop contact patterns, consistent with Pol V activity at or near these loci contributing to their chromatin structural organization. Color scale indicates normalized contact frequency difference (−0.80 to 0.10). **(C)** Metaplot enrichment analysis of percent 5-Methylcytosin DNA methylation (left y-axis, %5mC) and transcript abundance (right y-axis, log2(CPM+1)) over an 8 Kb window (±4 Kb) centered on the subset of 324 LCD borders that overlap high-confidence H3K27me3 ChIP-seq IDR peaks (IDR < 0.05), shown for WT (top) and *nrpe1* (bottom) under Control, High CO_2_, and pHigh conditions. CG (red), CHG (blue), CHH (green), and CNN (gold; all cytosine contexts, exclusive of other named contexts) methylation profiles are shown alongside mean transcript abundance (black). In WT Control plants, H3K27me3-marked LCD borders show a local depletion of transcript abundance at the border center flanked by elevated methylation, consistent with transcriptional silencing at PcG-marked domain boundaries. In High CO_2_ and pHigh plants, transcript abundance at these borders is attenuated relative to Control in WT, with the pattern altered in *nrpe1*, consistent with CO_2_-associated and Pol V-dependent modulation of transcriptional activity at H3K27me3-marked LCD borders. The absence of CHG/CHH methylation changes in *nrpe1* pHigh, compared with WT pHigh, is consistent with a role for Pol V-directed RdDM in maintaining non-CG methylation at these boundary loci across generations.

In WT pHigh, the aggregate LAL signal recovered but anchors more frequently contacted alternate partners further downstream of the LCD (Fig. 4A). Per-replicate observed/expected (O/E) scores confirmed this recovery, with WT pHigh means exceeding WT High across all biological replicates (Supplementary Fig. S5A-B). LALs were largely absent with the genetic ablation of Pol V (*nrpe1* Control), gained in *nrpe1* High, and persisted in *nrpe1* pHigh (Fig. 4A). Consistent with this, per-locus contact ratios showed that only 47.2% of LALs were attenuated in *nrpe1* High relative to *nrpe1* Control (below the 50% threshold) indicating a net LAL contact frequency gain. The reduced LAL signal in *nrpe1* Control resembled that of WT High (Fig. 4A), consistent with a reduction in Pol V targeting at LCD borders under elevated CO_2_. The LAL profile of WT Control resembled that of *nrpe1* pHigh, consistent with a contribution of RdDM to the formation and maintenance of LALs, LCDs, and local chromatin structure.

After observing a widespread CO_2_-dependent disruption of LCD- and LAL-associated chromatin contacts, we asked how elevated CO_2_ relates to the configuration of putative gene self-loops. Prior work established that chromatin along genes associated with transcriptional memory forms self-loops neighboring highly transcribed gene clusters, likely representing structural changes resulting from the binding of RNA polymerase transcription machinery (Lainé et al., 2009; Liu et al., 2016). We used a previously published set of 3,753 gene self-loop annotations (Liu et al., 2016) to explore the possibility that gene self-loop associated transcriptional memory could be disrupted under elevated CO_2_ conditions. The aggregate profiles of gene loop interactions reveal increased variability in interacting partners following elevated CO_2_ treatment, which was most pronounced in pHigh and further increased with the loss of Pol V (*nrpe1*) (Fig. 4B), consistent with altered chromatin loop configurations with reduced Pol V activity.

We analyzed levels of transcription and methylation in the CG, CHG, CHH, and CNN sequence contexts over all LCD borders (where H = A, C, or T and N = A, C, G, or T; CNN therefore denotes all cytosines pooled across sequence contexts) (Supplementary Fig. S6A). Elevated CO_2_ is associated with decreased transcript abundance upstream of H3K27me3-marked LCD borders in WT pHigh and *nrpe1* pHigh (Fig. 4C). Corresponding changes in DNA methylation in any sequence context over H3K27me3-marked LCD borders were near-undetectable (Fig. 4C; Supplementary Fig. S6B,C), an observation consistent with differential PcG activity contributing to transcriptional variation at LCDs independently of local 5mC changes. These results are consistent with LCDs and LALs participating in the plant response to elevated CO_2_ through altered chromatin loop contacts and transcript abundance near H3K27me3-marked border loci. The resemblance between *nrpe1* Control and WT High LAL profiles, supported by hundreds of loci, is consistent with elevated CO_2_ impairing Pol V targeting near LCD borders. The absence of transcriptional attenuation in *cmt2/3* pHigh (Supplementary Fig. S6A-C) is consistent with CMT2/3 activity maintaining silencing at these borders across generations.

### Elevated CO_2_ is associated with broad changes in gene and TE transcript abundance

Chromatin properties such as DNA methylation and histone modifications contribute to cellular responses and resilience in plant development, including epigenetic responses to abiotic stress (Cavalli & Heard, 2019; Lämke & Bäurle, 2017). To examine transcriptional changes associated with elevated CO_2_, we analyzed transcriptome data. We identified differentially abundant gene and TE (DAG, DATE) fragments in High and pHigh relative to paired Control plant transcriptomes, and we analyzed up- and down-regulated loci separately and together, because transcripts depleted or enriched in opposite directions may differ by spatial and epigenetic associations. Mapping DAG and DATE features to constitutive heterochromatin and euchromatin across all somatic chromosomes revealed genome-wide patterns of differential gene and TE abundance (Fig. 5A). As expected, heterochromatic gene and TE activation was prominent in *ddm1* plants. DDM1 contributes to epigenetic inheritance in dense chromatin regions together with the VARIANT IN METHYLATION (VIM) family of METHYLTRANSFERASE 1 (MET1) cofactors and H2A.W deposition (Lee et al., 2023). Loss of DDM1 (*ddm1*) alone caused heterochromatic DAG activation across somatic chromosomes independent of CO_2_ treatment, except for chromosome 2, where strong heterochromatic DAG activation was instead specific to WT, *cmt2/3*, and *nrpe1* pHigh (Fig. 5A, vertical orange box). The pericentromeric chromosome 2 DAGs are notable among CO_2_-responsive loci in being embedded within constitutive heterochromatin and showing high normalized expression specific to the pHigh generation, a pattern not recapitulated by DDM1 loss alone. Both *nrpe1* and *nrpd1* pHigh showed activation of DAGs and DATEs that emerged in the progeny generation (pHigh) rather than in the directly CO_2_-treated generation (High), a pattern consistent with a role for RdDM in chromatin-level silencing (Fig. 5A, horizontal orange boxes). The DAG and DATE activation in WT pHigh was broadly concordant with that of *nrpd1* and *nrpe1*, consistent with involvement of these core RdDM components in the response to elevated CO_2_ (Panda et al., 2023).

**Figure 5.**
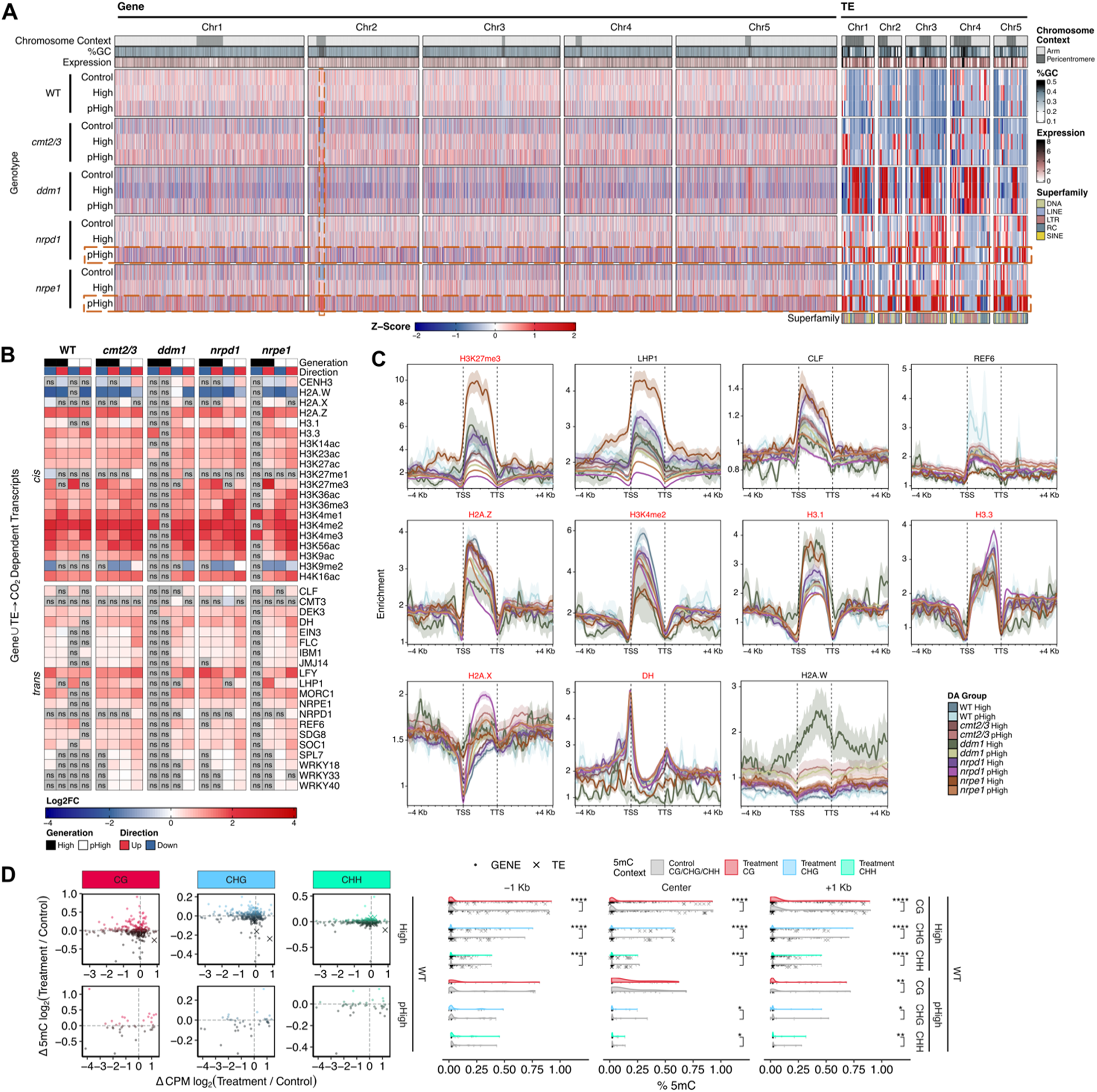
Elevated CO_2_ is associated with epigenetic reprogramming of genes and transposable elements across genetic backgrounds. **(A)** Genome-wide distribution of differentially abundant genes (DAGs, left) and TE fragments (DATEs, right) across all five somatic chromosomes in all genotypes and CO_2_ conditions relative to paired internal controls. Rows represent genotype-condition combinations; columns represent the ordered genomic position binned at chromosome scale. Color scale of the main heatmap reflects the Z-score of log2(CPM+1) normalized transcript abundance (blue to red), where the color intensity indicates the number of standard deviations away from the averaged ‘Expression’ track (rouge). ‘%GC’ indicates the fraction CpG content across the genome-wide distributions of DAG and DATE loci (light blue). Orange boxes highlight regions of delayed-onset activation (horizontal) and pericentromeric enrichment on chromosome 2 (vertical). Superfamily annotations specific to TEs are indicated below TE fragments on the right, showing enrichment of pericentromeric LTR and LINE DATEs. **(B)** Log2 fold-change heatmap of length-weighted mean histone ChIP-seq signal (1X normalized coverage) over DAG and DATE loci relative to all genes and TEs as a genomic background, stratified by direction of differential expression (Up, Down) across genotypes and CO_2_ conditions. Rows represent histone modifications and variants; non-significant values (Bonferroni-adjusted p > 0.05, Wilcoxon rank-sum test) are indicated as ‘ns’. **(C)** Weighted mean metaplots of statistically significant enrichments over a window spanning −4 Kb to +4 Kb relative to the TSS and TTS of statistically significant DAG and DATE loci, across all genotype-conditions (color-coded as indicated in the legend ‘DA Group’). Panels show H3K27me3, LHP1, CLF, REF6, H2A.Z, H2A.W, and additional factors associated with gene self-looping (Liu et al., 2016). Shaded regions indicate 95% confidence intervals. All ChIP-seq datasets used in panel C were generated under standard growth conditions from external sources and reflect general chromatin state associations rather than CO_2_-dependent changes in factor occupancy. **(D)** Left: MA plots showing the relationship between changes in log-transformed fractional 5-Methylcytosin levels (Δlog2(%5mC)) and log2 fold-change in transcript abundance for DAGs (circles) and DATEs (crosses) in CG, CHG, and CHH methylation contexts in WT High and pHigh plants relative to Control. Right: Beeswarm distributions of log-transformed fractional 5mC levels at DAG and DATE loci in the −1 Kb promoter region, gene body (Center), and +1 Kb downstream region across Control and Treatment conditions, stratified by sequence context. Significant differences between conditions are indicated (Wilcoxon rank-sum test; *** p < 0.001; ** p < 0.01; * p < 0.05; ns = not significant; FDR-corrected).

We next examined the enrichment of DAGs and DATEs for candidate factors associated with chromatin structure and the plant stress response, using published ChIP-seq datasets of mature plants grown under normal conditions (Fig. 5B-C). Because H3K27me3 was associated with LAL structure, particularly when Pol V is deficient, we asked whether its published binding sites also coincide with CO_2_-associated transcript abundance. H3K27me3 sites were enriched over CO_2_-associated DAGs and DATEs in all genotypes, and most strongly in *nrpe1* mutants (Fig. 5B). In an unperturbed genetic background (WT), down-regulated transcripts showed significant enrichment at LHP1-associated loci in both High and pHigh plants. In the absence of Pol V (*nrpe1*), only High up-regulated transcripts showed strong significant enrichment, with no directional preference of differentially abundant transcripts in *nrpe1* pHigh at LHP1-associated loci, consistent with a role for Pol V in establishing heritable CO_2_-dependent transcriptional down-regulation (Fig. 5B). The binding sites of the H3K27me3-modifying enzymes CURLY LEAF (CLF), LIKE-HETEROCHROMATIN PROTEIN 1 (LHP1), and RELATIVE OF EARLY FLOWERING 6 (REF6) were also significantly enriched (Fig. 5B). Metaplots in Fig. 5C visualize the average length-weighted mean spatial distributions of the statistically significant enrichments shown in Fig. 5B across the locus body. The observed enrichment of factors associated with gene looping (Liu et al., 2016) over CO_2_-associated DAGs and DATEs (Fig. 5B, 5C, red titles) is consistent with a role for local 3D chromatin folding in fine-tuning transcriptional activity. These patterns of enrichment align with a differential PRC2 activity model, consistent with mechanisms of imprinting and epigenetic inheritance (Heard & Martienssen, 2014). Establishing this precisely would require direct measurement with additional targeted enrichment assays.

Because the activities of enzymes central to maintenance DNA methylation and RdDM (CMT2/3, Pol V) were associated with changes in gene and TE abundance, we tested for differences in 5mC accumulation over DAG and DATE loci. WT DATEs showed increased abundance accompanying loss of 5mC in all sequence contexts (Fig. 5D). The global chromatin density changes shown in Fig. 1 are consistent with CO_2_-dependent changes in DNA methylation extending beyond individual DAG and DATE loci into their broader chromatin context. In WT DAGs, the differences in 5mC corresponded to whether the change occurred over the transcriptional start site (TSS) or the locus body. We examined this relationship using the length-weighted mean enrichment of DNA methylation over DAG and DATE loci (Fig. 5D, Supplementary Fig. S7). In WT High, significant differences in DNA methylation were identified across all sequence contexts at putative promoter regions (−1 Kb), the locus (Center), and downstream adjacent regions (+1 Kb) (Fig. 5D). In WT pHigh, significant differences in DNA methylation persisted over the locus and downstream regions, consistent with continued silencing of DAGs and DATEs. These results are consistent with CO_2_-dependent changes in DNA methylation and chromatin organization shaping transcript abundance at loci embedded within the structurally plastic 3D chromatin landscape characterized in the preceding sections.

## Discussion

The 3D genome underlies developmental and environmental response programs. The combination of sequence composition, epigenetic modifications, and the enzymes that interact with them cooperatively govern the shape of the genome, which in return provides structural support, aiding the reinforcement or gradual depletion of deterministic epigenetic modifications. While environmental stress promotes changes in genome structure, elevated CO_2_ is not inherently maladaptive to the plant. However, plant growth at elevated CO_2_ may introduce unexpected consequences, altering the way plants interact with abiotic and biotic factors within the environment (Kobayashi et al., 2006). In evolutionarily distant plants, elevated CO_2_ produces an accelerated growth phenotype that is maintained in ambient CO_2_ progeny (Panda et al., 2023). Elevated CO_2_ increases plant size and canopy density, but without adapting agronomic practices this produces a microclimate that is more permissive to pathogens (Manning & Tiedemann, 1995). Long-term growth at elevated CO_2_ levels will impact agricultural output and the standing ecology of complex biosystems. Conversely, plants primed with chromatin states shaped by elevated CO_2_ exposure could be a vehicle for increased agricultural output and a synthetic “off” switch engineered to mitigate any potential trade-offs would be advantageous. These principles will be critical to our understanding of long-term plant adaptation and evolution in diverse growing conditions, as managing the trade-offs associated with plant growth in a CO_2_-rich atmosphere becomes increasingly important.

One key observation is that genome conformation is associated with the growth environment through changes at facultative and constitutive heterochromatin. Our genome-wide analyses identified broad reorganization that accompanies locus-specific changes in chromatin interactions across progressively finer scales of the structural hierarchy of the genome. Shifts in A–B compartmentation that were retained in the progeny generation are consistent with roles for DDM1 and Pol V in chromatin restructuring associated with the plasticity of transcriptional regulation. The combination of high CO_2_ and the loss of key chromatin-altering proteins is consistent with interplay between pathways associated with CO_2_-dependent changes in 3D chromatin conformation.

The divergent chromatin responses of *nrpd1* and *nrpe1*, two mutants disrupting distinct successive steps in the RdDM pathway, provide insights into the mechanistic specificity of the CO_2_-associated chromatin response. *nrpd1* phenocopies WT chromatin expansion under elevated CO_2_, while *nrpe1* does not, indicating that the two branches of the RdDM pathways are not functionally equivalent in this context. One interpretation is that elevated CO_2_ impairs

Pol V targeting at a subset of loci, potentially through changes in nuclear organization or chromatin accessibility at targeted loci, rather than upstream siRNA biogenesis, and that this branch-specific sensitivity is mechanistically linked to the observed chromatin expansion. The condensation observed in *nrpe1* and *ddm1* pHigh may reflect an alternative response in which heterochromatin typically maintained by Pol V targeting reorganizes when that targeting is impaired. Targeted measurement of Pol V occupancy under elevated CO_2_ would test this model directly.

Considering the role of nuclear matrix constituent proteins in regulating genome density (Grob et al., 2014) under stress conditions (Wang et al., 2023), our data are consistent with a model in which the 3D genome may facilitate epigenetic plasticity through chromatin architecture. The high-resolution Hi-C maps indicate that LCDs are prominent and widespread structures in the *Arabidopsis thaliana* 3D genome. The borders of these chromatin domains form over developmentally associated loci that anchor chromatin loops between genes and promoters or CREs. The correspondence between changes in 3D genome structure and Pol V-targeted loci is consistent with a role for small RNA generation and transposon silencing in shaping the structure of the plant 3D genome through LCDs and LALs. The regulatory properties of LCDs differed between domains within and outside constitutive heterochromatin (Supplementary Fig. S4). LCD borders outside constitutive heterochromatin are consistent with a transcriptionally poised chromatin state neighboring highly transcribed clusters of loci. At high resolution, the differential chromatin looping at LCD borders that anchor H3K27me3-enriched LALs is consistent with a collaboration between components of RdDM- and PcG-mediated silencing in maintaining transcriptional programming (Hure et al., 2025). One possibility, which our data do not test directly, is that CO_2_-dependent methylation signatures directed by Pol V inform the placement of H3K27me3, such that reduced Pol V activity associated with elevated CO_2_ may modify the rate or penetrance of H3K27me3 deposition. In the context of the reported role of H3K27me3 and chromatin loops in facilitating promoter–promoter interactions (Liu et al., 2016) and in regulating the distance between CREs and their target loci, our findings extend candidate mechanisms by which plants may initiate heritable changes in local chromatin architecture and transcriptional state.

The consistent over-representation of pericentromeric DAGs on chromosome 2 across WT pHigh, *cmt2/3* pHigh, and *nrpe1* pHigh, but not in CO_2_-exposed plants (High) or in *ddm1*, warrants specific consideration. The pericentromeric region of chromosome 2 harbors a large nuclear mitochondrial DNA insertion (NuMT), a genomic region with a distinct sequence composition and chromatin properties relative to the surrounding pericentromere. The restriction of this DAG activation to the progeny generation is consistent with a process dependent on heritable chromatin state changes established during germination or early development rather than an acute CO_2_ response. Whether the chromosome 2 pericentromeric DAGs specifically overlap NuMT coordinates, and whether their DNA methylation profiles differ significantly from DAGs at other pericentromeric loci, are the most direct tests of this hypothesis. The primary observation that chromosome 2 pericentromeric loci are reproducibly activated in the progeny generation across three independent genotype comparisons is notable, warranting further investigation in future studies.

The observations reported here establish CO_2_-associated reorganization of the *Arabidopsis* 3D genome as a multilayered phenomenon, operating from the scale of chromosome-wide compartmentalization to kilobase-resolution chromatin loop domains and transcriptional reprogramming. The convergence of chromatin architectural change with locus-specific alterations in DNA methylation and transcript abundance, and the heritability of these states, positions 3D genome reorganization as a candidate mechanism through which plants encode and transmit chromatin states associated with environmental CO_2_ exposure. The *nrpe1* mutant, which enhances the heritability of the accelerated growth phenotype (Panda et al., 2023) alongside the strongest chromatin expansion and compartment attenuation in pHigh, identifies Pol V activity as a key modulator of this process. Future work resolving the causal relationships among chromatin accessibility, Pol V occupancy, H3K27me3 deposition, and chromatin loop structure under elevated CO_2_ will be essential for understanding how plants integrate environmental signals into heritable chromatin states.

## Methods

### Plant growth and genotypes

The 1,000 ppm [CO_2_] used in our high CO_2_ treatment was determined by doubling our readings of atmospheric CO_2_, to match predicted atmospheric CO_2_ levels reported previously (Riahi et al., 2011). The experimental design and replication scheme for tissue generation were as described in a previous study by the authors (Panda et al., 2023). Plants were grown at ambient CO_2_, High CO_2_ fixed at 1,000 ppm across two growth rounds, and the self-fertilized progeny of High CO_2_ plants were grown at ambient CO_2_ by seed descent in a third growth round. All A. thaliana (L.) Heynh were of the Columbia (Col) ecotype and grown at 22°C on Pro-Mix FPX soil in Conviron PGC Flex growth chambers under long-day conditions (16 h : 8 h, light : dark) with 200 μmol m−2 s−1 light. Wild type (WT) Col plants were grown in each condition, in each round of growth, and were used to model batch effects (described below) of different growth rounds and chambers. Mutant alleles used were as follows: *cmt2-7/cmt3-11t*, *ddm1-2*, *nrpd1* (SALK_128428), *nrpe1* (SALK_017795). For each genotype/condition, four flats of 24 plants were grown. Replicates and genotypes were positioned randomly within two independent growth chambers across three growth rounds. Three of four flats underwent destructive tissue collection: above-ground whole leaves were collected 25 days after planting from both ambient and high CO_2_ grown plants. The fourth flat remained in the growth chamber for propagation, and self-fertilized seeds were collected.

### Chromatin Conformation Capture (Hi-C)

Whole leaf tissue was used as input for the Phase Genomics Proximo™ Hi-C kit (Seattle, WA) for plants using Sau3AI restriction enzyme digestion. Libraries were shallow sequenced and checked for long-range links using the hic_qc software (github.com/phasegenomics/hic_qc) before sequencing on an Illumina NovaSeq6000 with S4 XP PE150 kits, targeting at least 100X raw coverage per individual. Raw Hi-C reads were aligned to the TAIR10.1 reference genome with Juicer v1.6 with default options to produce .hic contact matrices filtered for aligned reads with a MAPQ score ≥ 30. ENCODE statistics were aggregated from the standard output of the Juicer processing pipeline, which validated all Hi-C libraries generated in this study as having met or exceeded the optimal Hi-C library quality metrics defined by the ENCODE consortia (Supplementary Data S1). Pairwise correlations between all Hi-C libraries were produced using the weighted average of Stratum Adjusted Pearson’s Correlation Coefficients (SCC), a methodologically robust method for directly comparing Hi-C libraries at the replicate level (Yang et al., 2017). Modeling the genotype-by-treatment interactions at the replicate-level via SCC, we observed high reproducibility between replicate libraries (Supplementary Fig. S8B), therefore we merged them to obtain high resolution chromatin contact maps that achieved sub-kilobase resolution (Supplementary Fig. S8A). We resolved chromatin contact matrices into 1 Kb, 2 Kb, 5 Kb, 10 Kb, 20 Kb, 25 Kb, 50 Kb, and 100 Kb bins and normalized for distance dependent coverage biases using the Knight-Ruiz (KR) matrix balancing method (Knight & Ruiz, 2013).

Sau3AI digestion is blocked by cytosine methylation at its recognition site, posing the question of whether differential methylation between Hi-C libraries could be a source of variance confounding the interpretation of differential chromatin contacts. Matched EM-seq data were used to classify the methylation status of all Sau3AI digestion sites in each genotype and conditions with a minimum 5X coverage and a 10% 5mC threshold. Comparative analyses were supported predominantly by the set of restriction fragments whose flanking sites were unmethylated across all samples, representing a methylation invariant fragment set. Of 435,342 total Sau3AI fragments, 341,374 (78.4%) met this conservative criterion, covering 78.1% of the *Arabidopsis* genome. Contact frequency fold-change values computed on the invariant subset of fragments were highly concordant with those from the full, unfiltered fragment set (Pearson r > 0.9999, Spearman ρ > 0.9999, n = 341,374; KS statistic D = 0.175, p < 0.001; Supplementary Fig. S9), confirming that the architectural differences reported are not driven by differential cut-site recovery. The significant KS statistic indicates that the excluded fragments are compositionally distinct from the invariant set, confirming that the filter was selective rather than a random subsampling of the full fragment population.

The matrices produced using Juicer were visualized in Juicebox (Durand et al., 2016) and HiGlass (Kerpedjiev et al., 2018) for initial inspection and used to produce genome-wide 2D heatmap visualizations. The R package GENOVA (van der Weide et al., 2021) was used to produce the eigenvectors of auto-correlated observed over expected transformed Hi-C contact matrices and quantify compartment strength, calculated as the frequency of intra-compartment interactions (AA, BB) over the frequency of inter-compartmental interactions (AB, BA). The signed PC3 eigenvector of all libraries was computed independently for each chromosomal arm and DNase I Hypersensitivity (DHS) was used to orient positively signed intervals with open chromatin. By this method, the third principal component (PC3) eigenvector best captured chromatin compartmentalization and was most concordant with previous studies (Wang et al., 2023). PC1 and PC2 were dominated by the chromosome-arm/centromere axis, whereas PC3 showed the strongest correlation with previously published *Arabidopsis* A/B compartment annotations (Fig. S3). Therefore, the signed PC3 eigenvector was used to annotate A and B compartments at 20 Kb resolution, designating consecutive positively or negatively signed 20 Kb bins as A or B compartments, respectively. Compartment switching in High and pHigh treatments relative to paired controls was determined using bedtools (Quinlan & Hall, 2010) intersect function, constrained by a 50% minimum positional overlap of the query compartments with reference Control compartments.

The annotation of high-confidence LCDs was accomplished using the hicFindTads function of the HicExplorer (Wolff et al., 2020) software applied to normalized matrices binned to 1 Kb resolution (parameters: –minDepth 3000, –maxDepth 30000, –threshold 0.05, –delta 0.01, – correctForMultipleTesting fdr). Aggregate analysis of LCDs interaction frequencies were produced using the CoolPup software (Flyamer et al., 2020), and visualized with Matplotlib. For LCD-Associated Loop (LAL) pile-ups, contacts were aggregated and rescaled to 87 × 87 pixel windows, centered on the central 1 Kb bin of each LCD with flanking regions extending ±50 Kb on each side, such that one loop anchor falls within the flank. LCD coordinates were reoriented to a common 5ʹ to 3ʹ convention (start-to-end) prior to aggregation so that loops are summed in a consistent orientation, resulting in the asymmetric pattern of the resulting pile-up. Histone-mark and variant enrichment profiles over LCD borders were grouped into clusters (C1–C5) by k-means clustering, where k produces biologically meaningful separation of the per-border mark-enrichment vectors. All other Hi-C matrix heatmaps were visualized using FanC (Kruse et al., 2020) with custom Matplotlib and Seaborn (Waskom, 2021) functionalities.

Differences in chromatin interaction decay between conditions were assess using an Ordinary Least Squares (OLS) linear model of the form: Contacts ∼ condition + batch + distance, where “condition” is the CO_2_ treatment (High or pHigh vs. paired Control), “batch” is the biological growth round included as a fixed blocking factor, and “distance” is the pairwise genomic separation included as a continuous covariate. For WT pHigh comparisons, a paired “pControl” replicate set was used as the reference in place of the standard ambient CO_2_ Control. The p-value for the condition term was extracted from the fitted model for each genotype-condition comparison. Where OLS model fitting failed, a Welch’s t-test was used a conventional fallback. P-values from the High and pHigh comparisons within each genotype were corrected for multiple testing using the Benjamini-Hochberg false discovery rate (FDR) method. Significant comparisons (FDR < 0.05) are indicated within Fig. 1C and color-coded by condition.

LCD borders were tested for enrichment of TE loci occupies by Pol V (NRPE1; SRP014189) and LHP1 (SRP068984) using a statistical permutation-based approach. All external ChIP-seq datasets used for LCD border enrichment analysis and downstream histone mark enrichment analyses were generated from mature *Arabidopsis* arial tissue grown under standard conditions, reflecting the baseline chromatin state of relevant loci rather than CO_2_-dependent changes in epigenetic modification deposition or transcription factor occupancy that were directly measured. For each transcription factor, the target set was defined as the intersection of Araport11-annotated TE coordinates with IDR-filtered ChIP-seq peaks from the respective dataset. The observed fraction of LCD borders overlapping target TE loci was compared against a null distribution generated by 1,000 random shuffles of LCD border coordinates using bedtools shuffle (parameters: -chrom, -noOverlapping), applied to the LCD borders flanked by ±4 Kb using bedtools slop for consistency with other overlap analyses throughout the manuscript. Empirical p-values were computed as the proportion of shuffles yielding an overlap fraction at or above the observed value. Co-occurrence of Pol V- and LHP1-enriched borders sets was assessed using Fisher’s exact test on a 2×2 contingency table of border membership.

Chromatin loop anchors in WT Control were identified using hicDetectLoops from the HicExplorer software (Wolff et al., 2020) applied to the merged WT Control Hi-C contact matrix binned to 1 Kb resolution (parameters: –windowSize 5, –peakWidth 2, –maxLoopDistance 30,000, –pValuePreselection 0.05, –pValue 0.05). Left and right anchor coordinates were extracted from the resulting BEDPE file, combined, sorted, and merged using bedtools merge to produce non-redundant loop anchor regions. LCD borders were tested for overlap with loop anchors by applying a ±4 Kb flank to the LCD border coordinates using bedtools slop and intersecting with the merged anchor set using bedtools intersect (-u). The co-occurrence of H3K27me3 and H2A.Z IDR peaks at LCD borders was assessed by sequential bedtools intersect commands of ±4 Kb-flanked LCD borders with H3K27me3, then H2A.Z IDR peak sets. The spatial definition for the subset of 324 LCD borders used specifically in the Fig. 4C analysis was a derived from a direct overlap of unflanked 1 Kb LCD border bins with H3K27me3 IDR peaks without extension.

To prepare the matrices for 3D modeling, a batch script was implemented in MATLAB (Ver. R2024a) to automatically convert .hic formatted matrices into chromosome-chromosome pairwise contact matrices saved as .txt files. The five *Arabidopsis* somatic chromosomes yielded 10 chromosome-chromosome pairwise matrices. Combined matrix blocks were loaded into full upper and lower sparse triangular matrices. Missing values and all-zero columns and rows were removed to produce a 5,920 × 5,920 contact matrix. The 3D models were constructed using Max3D (Oluwadare et al., 2018) by partially re-implementing the MATLAB algorithm to improve computational efficiency via parameter optimization. The models were saved as 3D coordinates corresponding to 20 Kb genomic bins annotated with chromosome information. Each 3D model is represented as a 5,920 × 3 matrix. A representative WT sample was selected as a reference and registered remaining 3D models to this reference model using Procrustes alignment with translation and rotation transformations. Each 3D model (5,920 × 3 matrix) was transformed into a long vector (17,760 × 1) for Principal Component Analysis (PCA). A 3D Gaussian kernel density estimate was computed from each 3D model with a choice of bandwidth and mesh grids. To visualize the density in 2D, the heatmap of the maximum projection of the density along the z-axis was plotted for all chromosomes.

### RNA sequencing (RNA-seq)

Total RNA was isolated from leaf tissue using Trizol prep followed by phenol chloroform isolation. Poly(A)+ mRNA was enriched using NEBNext mRNA poly(A) enrichment module and the library was generated using NEBNext Ultra II Directional RNA sequencing using the manufacturer’s protocol with the following exceptions: RNA fragmentation was performed for 7 minutes (reduced from 15 minutes), size selection bead enrichment was performed according to Appendix 6 in the protocol, and 13 PCR cycles were used for amplification. Libraries were pooled equimolarly before QC and sequencing on NovaSeq 6000 at Genomics Technology Core at University of Missouri, Columbia. Raw reads (Illumina PE150) were trimmed for adapters using Trimmomatic (Bolger et al., 2014) with the following parameters: ILLUMINACLIP:Trimmomatic-0.39/adapters/NEBNext-PE.fa:2:30:10:2:true MINLEN:30. Trimmed reads were then mapped to the *Arabidopsis* genome (TAIR10.1 annotation) using STAR (Dobin et al., 2013) (parameters: –outMultimapperOrder Random, –outSAMtype BAM SortedByCoordinate, –outFilterMultimapNmax 50, –outFilterMatchNmin 30, –alignIntronMax 10000, –outSAMattributes All).

Raw counts of genes and TEs were quantified with FeatureCounts (Liao et al., 2014) using ‘exon’ and ‘transposable_element’ annotations, respectively (Supplementary Fig. S10). To maximize the certainty of aligned read assignment to annotated TEs, only counts from TE fragments that have less than 20% overlap with genes were retained. Multifactorial differential expression analyses were performed using a Generalized Linear Model (GLM) for each genotype and treatment relative to paired internal control transcriptomes. Growth round and chamber metadata were incorporated as blocking factors to inform the model and control for batch effects. Genes and TEs with a residual p-value < 0.05, Log2 fold-change ≥ 1, and FDR < 0.05 were retained as significant differentially abundant RNA. Normalized abundances of DAGs and DATEs were generated using EdgeR (Chen et al., 2025) and exported for downstream analysis and visualization (Supplementary Fig. S11). To compare the profile of DAGs and DATEs across genotypes and CO_2_ treatment, we standardized their normalized expressions by the standard deviation (Z-Score) from the aggregate mean. All CO_2_-dependent *Arabidopsis* gene and TE accessions with respect to each genetic background in Supplementary Data S2.

### Chromatin Immuno-precipitation sequencing (ChIP-seq) and external data

External ChIP-seq data sets were downloaded from The Plant Chromatin State Database (Liu et al., 2018) and ChIP-Hub (Fu et al., 2022) quantifying enrichment normalized to 1x coverage. All external ChIP-seq datasets were generated from mature *Arabidopsis* plants that were grown under standard conditions and reflect the baseline chromatin state of relevant loci rather that CO_2_-dependent changes in deposition and occupancy. The Irreproducibility Discovery Rate (IDR) was used to ensure high certainty of peak enrichment. Only peaks with an IDR score below 540, calculated as int(−125log2(0.05)), were used, as these have a less than 5% probability of being false positives. The overlap of LCD borders with IDR peaks was identified using bedtools intersect (Quinlan & Hall, 2010). Normalized ChIP-seq signals were mapped to DAG and DATE loci using BEDOPS bedmap wmean (Neph et al., 2012). Enrichment of epigenetic modifications over significant DAG and DATE loci were visualized using the ComplexHeatmap R package. Gene self-loops (n=3,753) were obtained from a previous study and were annotated as described (Liu et al., 2016). The expression distribution data of self-looped genes across genotypes and conditions in this study are provided in (Supplementary Data S4).

### Enzymatic Methyl-seq (EM-seq)

Enzymatic methylation sequencing and quantifications were performed as previously described (Panda et al., 2023). The percent 5mC enrichment over LCD boundaries was calculated using the deepTools (Ramírez et al., 2014) computeMatrix function. Length-weighted mean methylation scores over gene and TE loci were calculated using BEDOPS bedmap wmean (Neph et al., 2012) to account for differences in length between mapped loci.

## Quantification and statistics

Statistical tests were performed using the non-parametric Wilcoxon rank sum test, unless stated otherwise. Statistically significant effects were assigned when *p* < 0.05 and FDR < 0.05.Enrichment of Pol V- and LHP1-bound TE loci at LCD borders was assessed via permutation (n =1000 shuffles; empirical p-values reported as described above). Co-occurrence of Pol V- and LHP1-enriched LCD border populations was assessed by Fisher’s Exact test.

## Supplementary Data

Supplementary Data S1 (separate file). Table of ENCODE mapping statistics for all Hi-C matrices in this study.

Supplementary Data S2 (separate file). Tables of CO_2_-dependent differentially abundant genes (DAGs) and TEs (DATES). The data include annotation information, and raw and normalized estimates of abundance of fragments in each genotype and treatment condition.

Supplementary Data S3 (separate file). Table summary of all external data used in this study complete with DOI references to the original studies.

Supplementary Data S4 (separate file). Table of directly measured self-looped gene and TE transcriptional abundance per all genetic backgrounds and treatment conditions featured in this study.

## Acknowledgments

We thank L. McClain (Donald Danforth Plant Science Center) and S. Kenney (Donald Danforth Plant Science Center) for assistance with plant growth. We thank M. Gehan (Donald Danforth Plant Science Center), N. Fahlgren (Donald Danforth Plant Science Center), S. Pandey (Donald Danforth Plant Science Center), and R.J. Schmitz (Department of Genetics, University of Georgia) for valuable scientific discussions and suggestions. This work was supported by the National Science Foundation (NSF) grant EF-1921724 (to B.C.M. and R.K.S.).

## Author Contributions

S.A.L., A.H., B.C.M., and R.K.S. conceptualized the research. S.A.L., M.L., K.P., and A.H. developed the methodology and performed the investigation. S.A.L. and M.L. performed the visualization and formal analysis. S.A.L. curated the data. B.C.M. and R.K.S. acquired funding, administered the project, and provided supervision. S.A.L. and B.C.M. wrote the original draft of the manuscript, and all authors (S.A.L., M.L., K.P., A.H., R.K.S., B.C.M.) contributed to the review and editing of the manuscript. All authors reviewed and approved the final version of the manuscript.

---

**Supplementary Figure S1.**
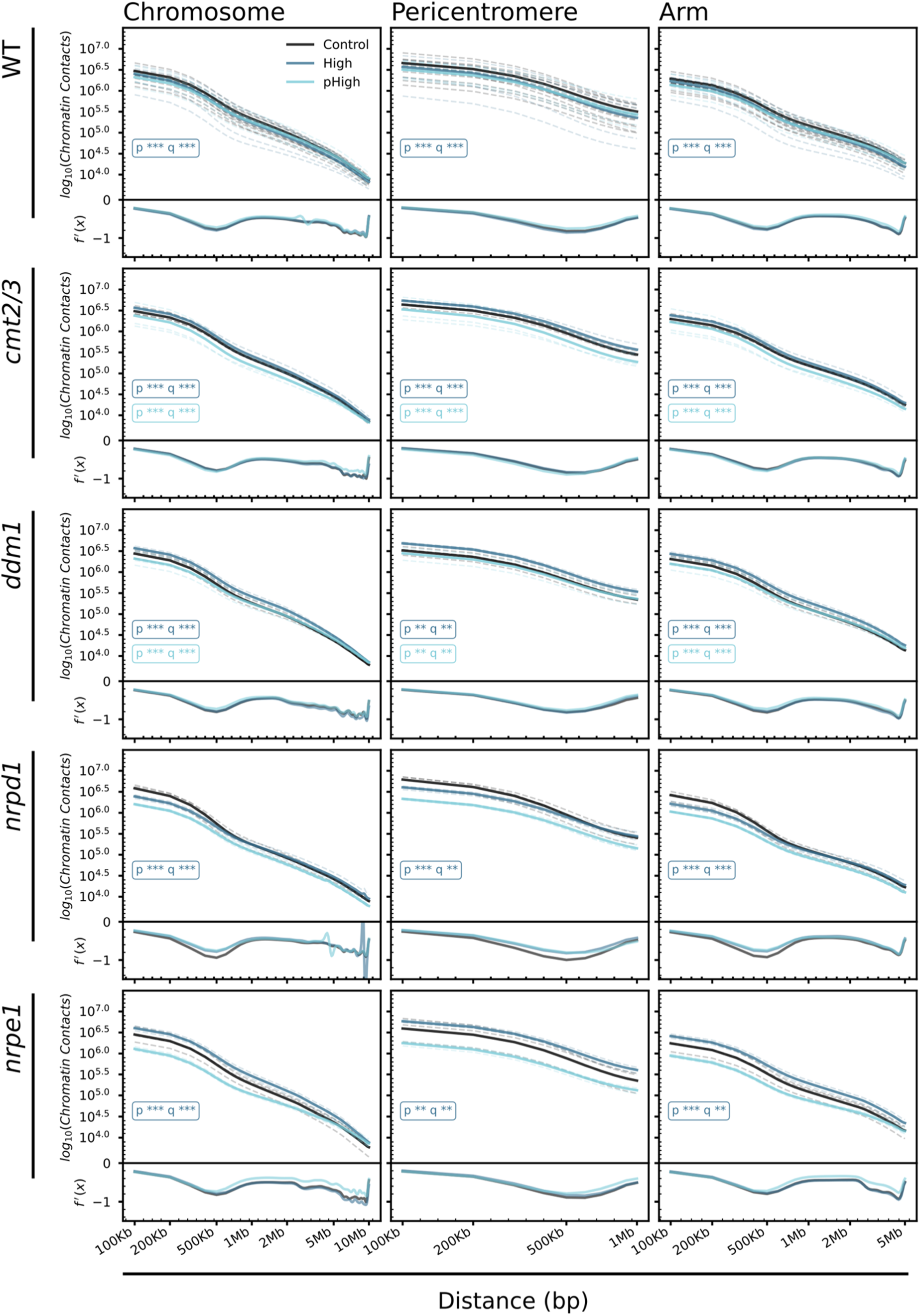
Replicate-resolved chromatin contact frequency decay curves across genotypes, CO_2_ conditions, and genomic contexts. Log10-transformed chromatin contact frequency as a function of genomic distance (bp, x-axis) for all five genotypes examined (WT, *cmt2/3*, *ddm1*, *nrpd1*, and *nrpe1*) (rows) stratified by genomic context: whole chromosome (left), pericentromeric heterochromatin (center), and chromosome arms (right). Contact frequency decay is shown for ambient CO_2_ (Control; black), elevated CO_2_ (High; dark blue), and progeny of High CO_2_ plants returned to ambient conditions (pHigh; light blue). Solid lines represent the non-redundant consensus profile across all biological replicates per treatment condition; dashed lines represent individual replicates, illustrating within-condition reproducibility. Below each decay plot, the Lagrange derivative residual f(x) trace shows the difference between High or pHigh and the paired Control decay profile, facilitating visualization of condition-dependent shifts in contact frequency at each distance scale. Statistical significance of differences between conditions is indicated within each panel (Wilcoxon rank-sum test; *** *P* < 0.001; ** *P* < 0.01; * *P* < 0.05; absent = not significant; FDR-corrected) using the full statistical power of individual replicates. In WT plants, the High CO_2_ condition is associated with a reduction in short- to mid-range contact frequency (approximately 100 Kb–1 Mb), consistent with chromatin decondensation, an effect that is most pronounced in the pericentromeric context. The *nrpd1* mutant shows a similar or enhanced decondensation pattern relative to WT, while *nrpe1*, *cmt2/3*, and *ddm1* show attenuated or biphasic patterns, consistent with distinct roles for the Pol IV and Pol V branches of the RdDM pathway and for DDM1-mediated chromatin remodeling in mediating CO_2_-associated changes in chromatin compaction. Contact matrices were produced as described in the Methods, subsampled to the lowest in-group depth, and normalized using the Knight-Ruiz (KR) balancing method (Knight & Ruiz, 2013) prior to all direct comparisons.

**Supplementary Figure S2.**
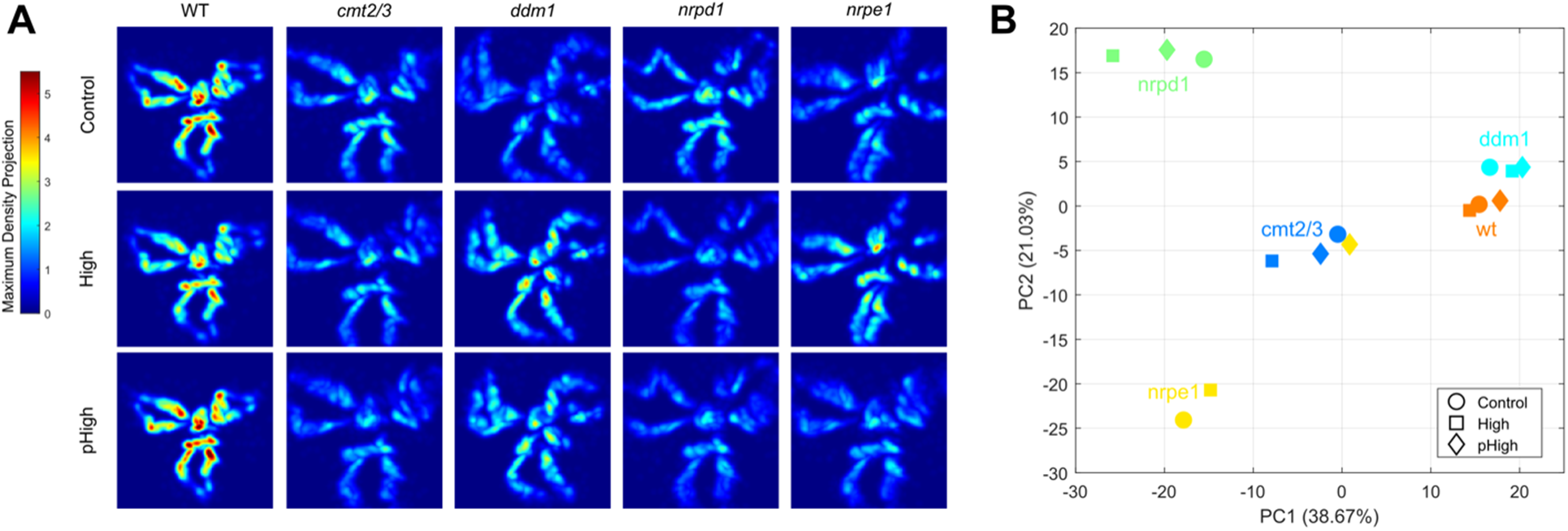
3D genome models reveal CO_2_-associated changes in chromatin density and spatial organization across genetic backgrounds. **(A)** Maximum density projection heatmaps derived from 3D genome models of all five somatic *Arabidopsis* chromosomes in all five genotypes (WT, *cmt2/3*, *ddm1*, *nrpd1*, *nrpe1*; columns) under Control, High CO_2_, and pHigh conditions (rows). Models were constructed from whole-genome Hi-C contact matrices at 20 Kb resolution using Max3D (Oluwadare et al., 2018) and registered to a common WT Control reference via Procrustes alignment. Color scale reflects maximum density projection intensity, where warmer colors indicate regions of greater chromatin compaction associated with centromeres and pericentromeric regions. CO_2_-associated chromatin expansion in WT High, and the expanded conformation with increased chromocenter density in WT pHigh, are consistent with the contact frequency decay profiles shown in Fig. 1C and Supplementary Fig. S1. Genotype-dependent differences in density distribution are visible across conditions, particularly in *ddm1*, which shows markedly altered spatial organization relative to WT under all conditions, consistent with DDM1’s role in maintaining constitutive heterochromatin compaction. **(B)** Principal component analysis (PCA) of 3D genome models across all genotypes and conditions. Each point represents a single merged Hi-C library 3D model; shapes indicate condition (circle = Control, square = High, triangle = pHigh) and colors indicate genotype as labeled. PC1 (38.97%) and PC2 (27.07%) together explain 66.04% of total variance in 3D genome configuration. Points cluster primarily by genotype along PC1, with CO_2_ treatment producing condition-dependent displacement along PC2 within each genotype, consistent with CO_2_-associated changes in global chromatin architecture being secondary to genotype-driven differences in 3D genome organization.

**Supplementary Figure S3.**
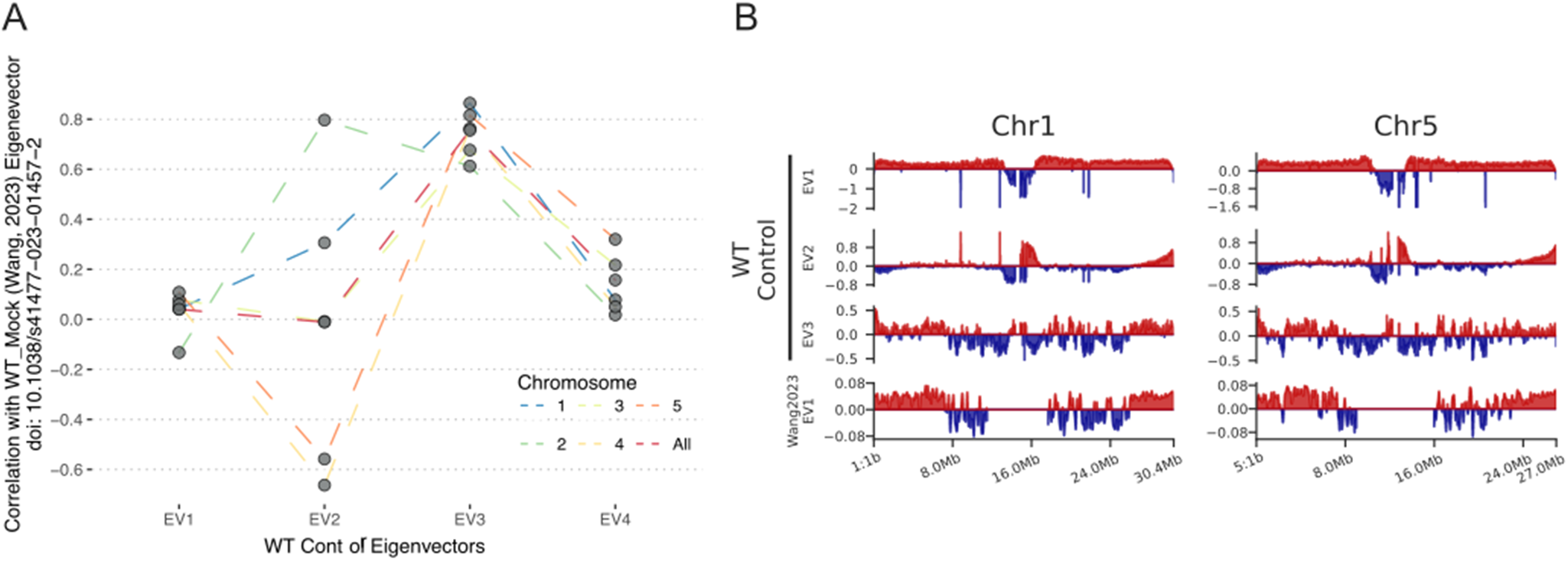
Validation of PC3 (EV3) for A/B chromatin compartment annotation in *Arabidopsis*. **(A)** Pearson’s *r* correlation coefficients between WT Control eigenvectors EV1– EV4 and the published chromatin compartment eigenvector (EV1) WT mock-treated Arabidopsis (Wang et al., 2023). Colored lines connect per-chromosome correlation values (chromosomes 1–5 and genome-wide); individual data points represent the observed correlation (Pearson’s *r*) for each chromosome context. EV3 shows substantially higher correlation with the published reference compartment eigenvector across all five somatic chromosomes than EV1, EV2, or EV4, supporting its use for A/B compartment annotation. **(B)** Genome-wide eigenvector profiles for WT Control EV1, EV2, and EV3, and the EV1 reference (Wang et al., 2023), plotted along chromosomes 1 and 5. Positive and negative values are shown in red and blue, respectively. EV3 shows the closest spatial correspondence to the published reference, recapitulating the known pattern of A compartment (euchromatin, chromosome arms) and B compartment (pericentromeric heterochromatin) identity. EV1 and EV2 reflect orthogonal sources of variance in the contact matrix; primarily the separation of chromosome arms from centromeric heterochromatin, rather than compartment identity. Panels A and B support the use of PC3 for A/B compartment annotation in this dataset, consistent with previous reports in *A. thaliana* (Wang et al. 2023).

**Supplementary Figure S4.**
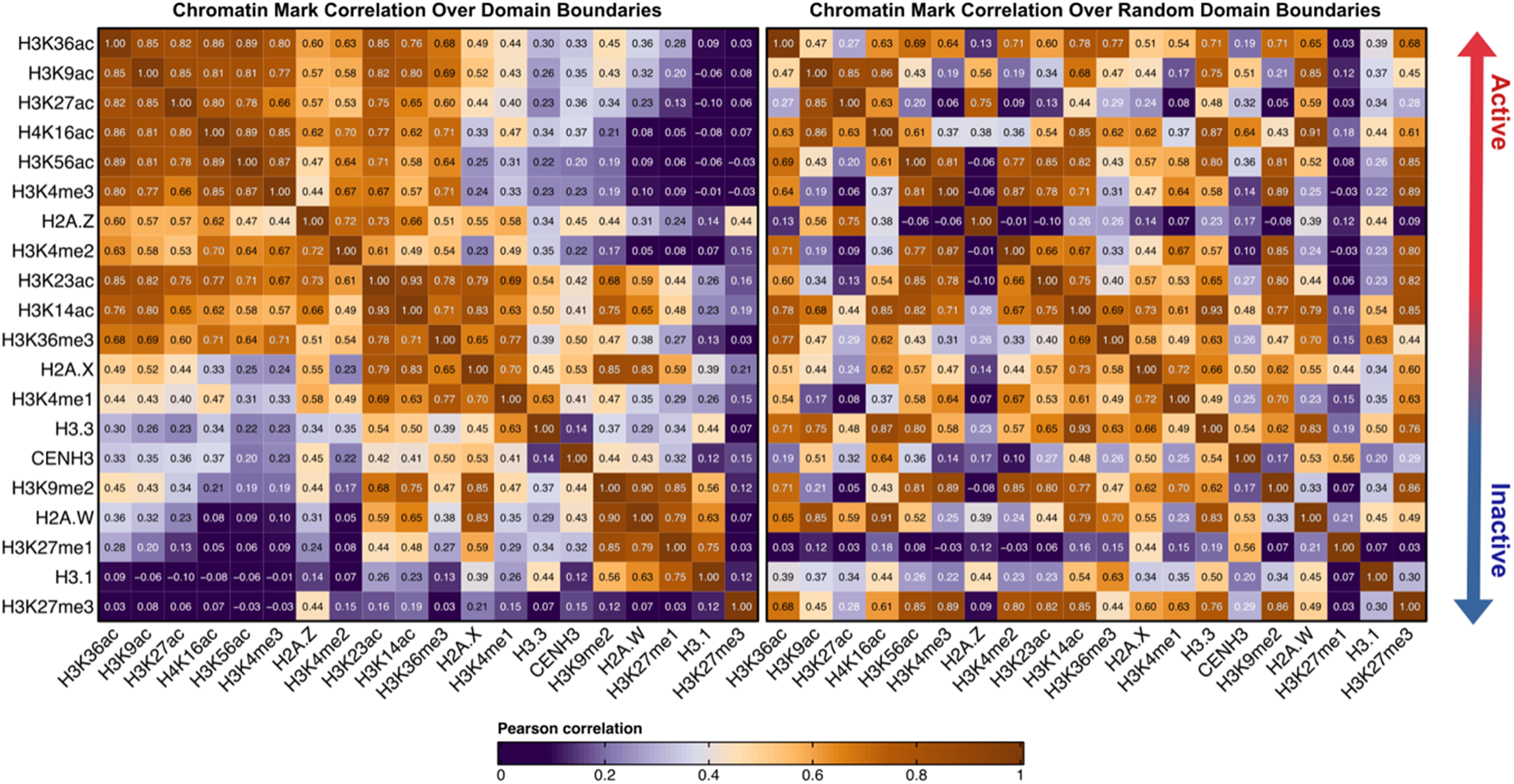
Chromatin mark co-enrichment at LCD borders reflects a structured epigenetic landscape distinct from random genomic loci. Pairwise Pearson correlation matrices of normalized ChIP-seq signal for 20 histone modifications and variants at annotated LCD borders (left) and an equal number of randomly positioned control regions (right), calculated using external ChIP-seq datasets (Fu et al., 2022; Liu et al., 2018). Marks are ordered by hierarchical clustering and annotated along the right axis by chromatin state association: active marks associated with transcriptionally active chromatin (red arrow, top) and associated with silent chromatin (blue arrow, bottom). Color scale indicates Pearson correlation coefficient (orange/warm = positive; purple/cool = negative or near-zero), with gradient boundaries in accordance with standard Cohen’s estimation of effect size. At LCD borders, active marks show strong positive co-correlations within the active cluster and negative correlations with inactive marks, indicating a structured bimodal chromatin landscape consistent with the k-means clustering presented in Fig. 3E. This structured co-enrichment pattern is not conserved within a background of randomly positioned control regions (right panel), where inter-mark correlations are weaker and less coherent, supporting the interpretation that LCD borders punctuate a non-random chromatin environment. H3K27me3 shows a moderate positive correlation with H2A.Z at LCD borders (Pearson r = 0.44), consistent with their co-enrichment at a subset of euchromatic LCD borders as shown in Fig. 3C–E.

**Supplementary Figure S5.**
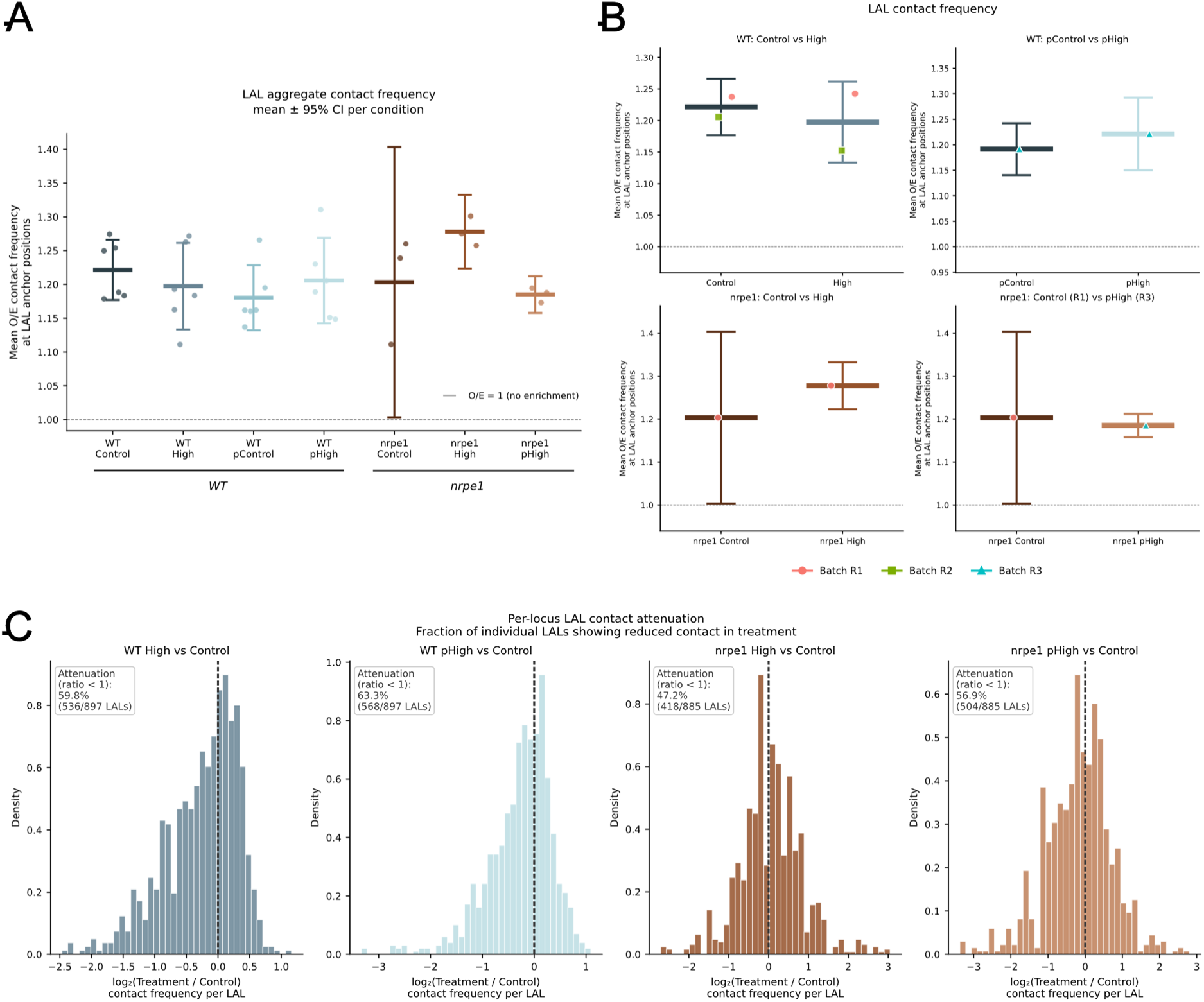
Quantitative validation of LAL contact attenuation across biological replicates. **(A)** Per-replicate aggregate observed/expected (O/E) contact frequency at LAL anchor positions for WT and *nrpe1* across all CO2 conditions. For each replicate, contacts were aggregated across all 749 LCD borders that anchor LALs using coolpup with KR balancing, local normalization with shifted controls (n = 10 shifts, 10–50 Kb), and rescaling to 87 × 87 pixel windows. The O/E score was extracted from the downstream loop anchor region (pixels 84–86 in both axes), where loop anchor contacts are concentrated in the oriented pile-up. Horizontal bars indicate the condition mean; error bars indicate the 95% confidence interval computed from the t-distribution across all replicates per condition. The reference line at O/E = 1.0 indicates no enrichment relative to stochastic background. **(B)** Batch-level mean O/E scores for the same conditions. Each point represents the mean O/E across replicates within a single biological growth round (batch), colored by batch identity. Horizontal bars indicate the overall condition mean; error bars indicate the 95% confidence interval across per-replicate scores. LAL contact frequency is modestly reduced in WT High relative to WT Control and recovers in WT pHigh. In nrpe1, LAL contact frequency increases in High relative to Control, consistent with LAL gain in this genotype as shown in Fig. 4A. The *nrpe1* cross-round panel (bottom right) compares nrpe1 Control (R1) with nrpe1 pHigh (R3); because these conditions derive from different growth rounds this panel is shown for descriptive completeness and does not constitute a formally paired comparison. **(C)** Per-locus LAL contact attenuation. For each of 897 unique loops anchored at 749 LAL borders (WT comparisons) or 885 loops (*nrpe1* comparisons), the contact frequency at the loop pixel was extracted from merged condition cooler files, normalized by total library contacts (RPM), and expressed as the log2 ratio of treatment to control. Distributions shifted left of zero indicate net contact attenuation in the treatment condition. The fraction of individual loops with a ratio below 1 is annotated for each panel. In WT, 59.8% and 63.3% of LALs show contact attenuation in High and pHigh, respectively. In nrpe1 High, 47.2% of loops show attenuation below the 50% threshold, indicating net contact gain relative to nrpe1 Control, consistent with the LAL gain observed in the aggregate pile-up (Fig. 4A). In *nrpe1* pHigh, 56.9% of LALs show attenuation relative to *nrpe1* Control.

**Supplementary Figure S6.**
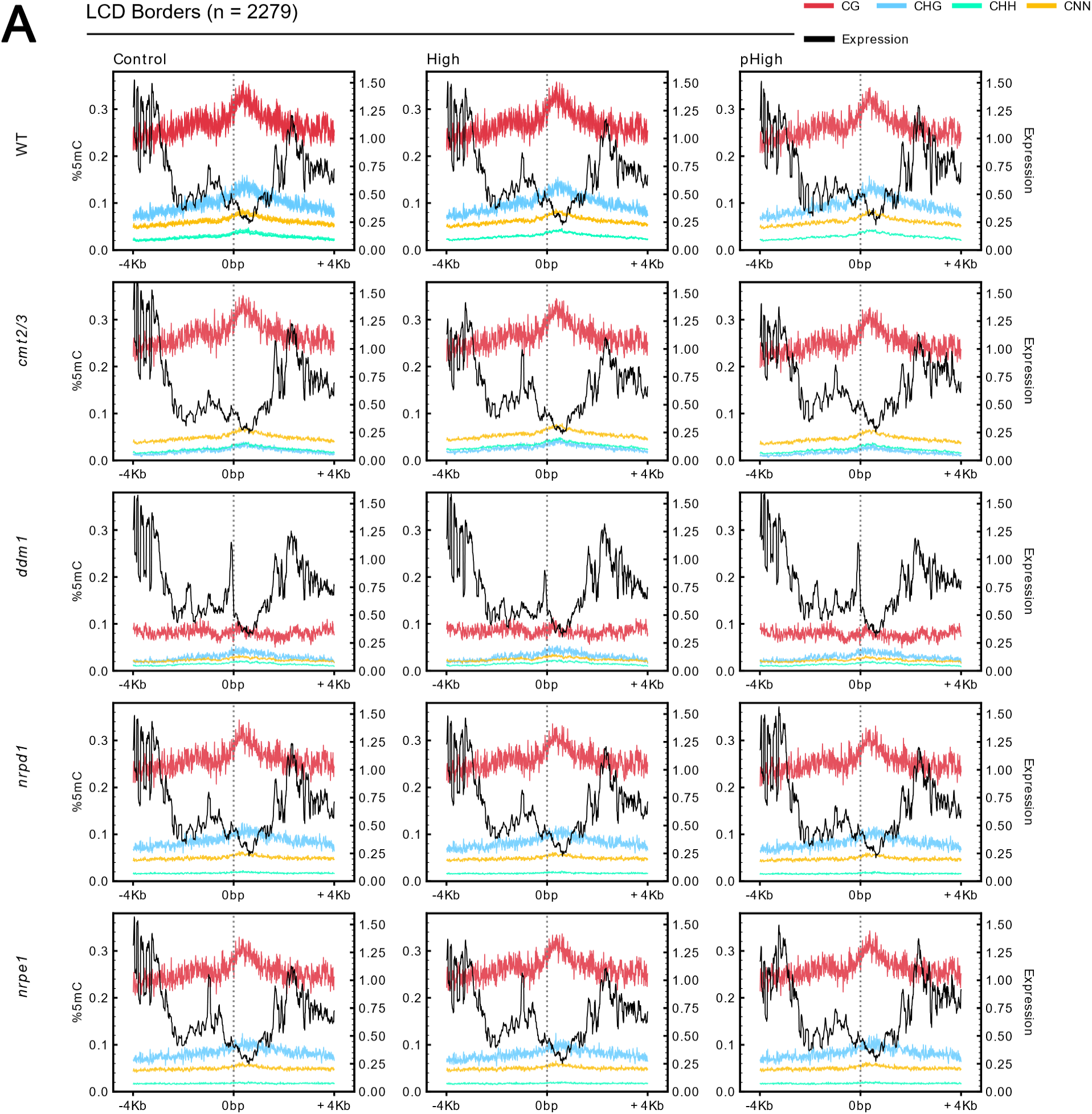
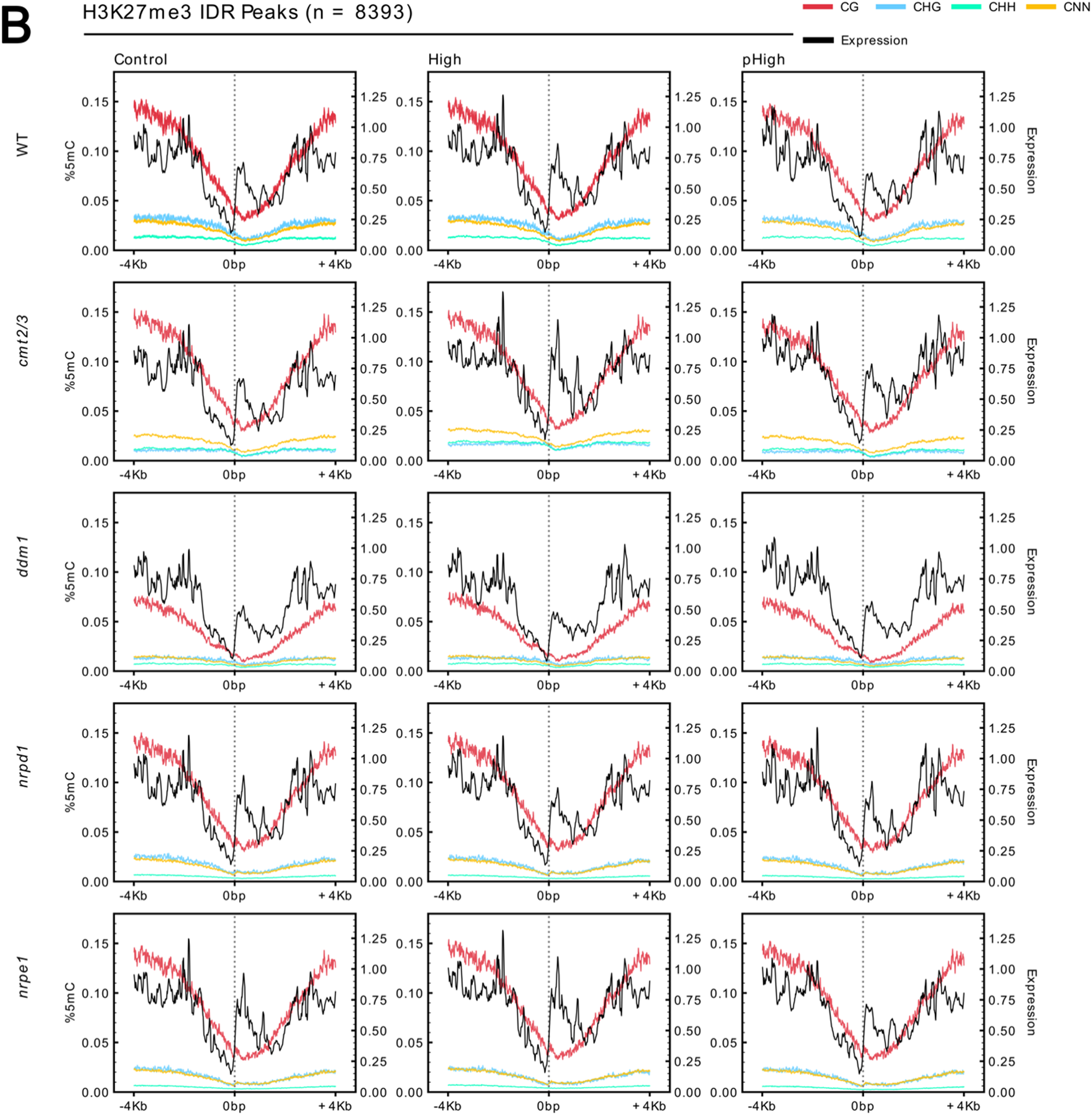
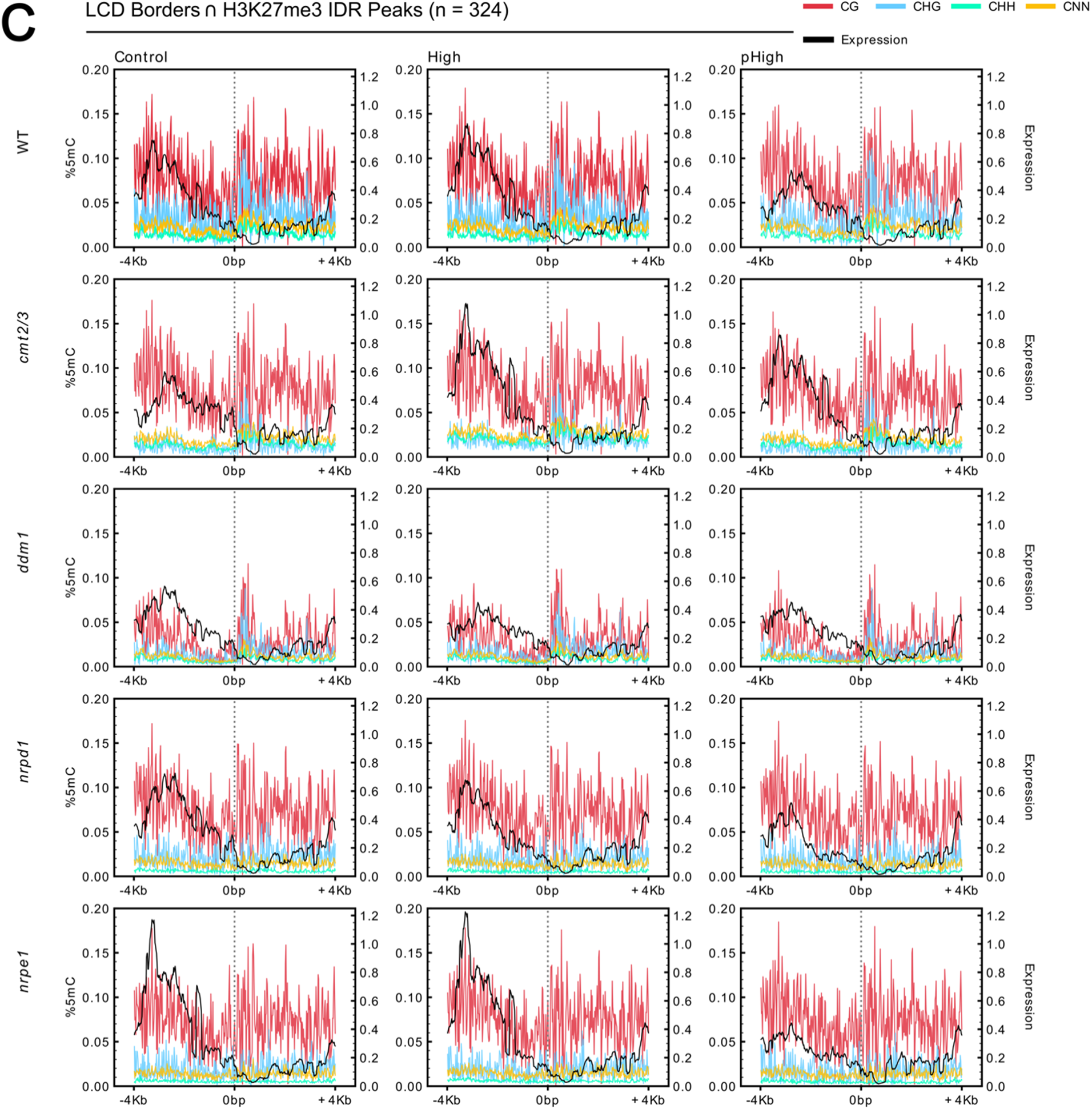
LCD borders are associated with distinct patterns of DNA methylation and transcriptional activity, particularly at H3K27me3-marked loci. Metaplot enrichment analyses of DNA methylation and transcript abundance over an 8 Kb window (±4 Kb) centered at 0 bp, across all five genotypes (WT, *cmt2/3*, *ddm1*, *nrpd1*, *nrpe1*; rows) and CO_2_ treatment conditions (Control, High, pHigh; columns). Left y-axes (% 5mC) indicate the length-weighted mean fractional 5-methylcytosine composition in CG (red), CHG (blue), CHH (cyan), and CNN (gold; all cytosines exclusive of other named contexts; H = C, A, or T; N = C, A, T, or G) sequence contexts. Right y-axes (Expression) indicate mean log2(CPM + 1) transcript abundance. Dashed vertical lines at 0 mark the center of LCD border regions in aggregate. Methylation enrichment was calculated using EM-seq data and transcript abundance from RNA-seq data, as described in the Methods. (A) Metaplot profiles centered on all 2,279 annotated LCD borders (n = 2,279). Expression shows a characteristic dip at the LCD border in WT Control plants, consistent with the transcriptionally poised or repressed chromatin state at domain boundaries. CG methylation is broadly elevated across the window in all genotypes, while CHG and CHH methylation show context- and genotype-dependent variation, particularly in *ddm1* and *cmt2/3* backgrounds. (B) Metaplot profiles centered on all high-confidence H3K27me3 ChIP-seq IDR peaks (n = 8,393; IDR < 0.05) from external datasets (Fu et al., 2022; Liu et al., 2018). H3K27me3-marked loci show a pronounced depletion of transcriptional activity at the peak center relative to flanking regions in WT Control plants, consistent with Polycomb group-mediated transcriptional silencing. CG methylation profiles are broadly similar to Panel A, while CHG and CHH methylation show reduced enrichment at H3K27me3 peak centers in WT, consistent with the known mutual exclusivity of RdDM-directed and PcG-directed silencing pathways at select loci. (C) Metaplot profiles centered on the subset of LCD borders that overlap high-confidence H3K27me3 IDR peaks (n = 324). This subset shows the most pronounced expression depletion at the border center of the three comparisons, consistent with H3K27me3-marked LCD borders representing a transcriptionally silent subset with co-enrichment of PcG-associated features. The attenuation of this expression depletion in High and pHigh conditions in WT, and the genotype-dependent differences in *nrpe1* and *cmt2/3* backgrounds, is consistent with CO_2_-associated and RdDM-dependent modulation of transcriptional activity at LCD borders frequented by PcG-associated machinery. Comparison across panels A–C supports the interpretation that H3K27me3 enrichment at LCD borders marks a functionally distinct chromatin state associated with transcriptional silencing and the CO_2_-dependent transcriptional response.

**Supplementary Figure S7.**
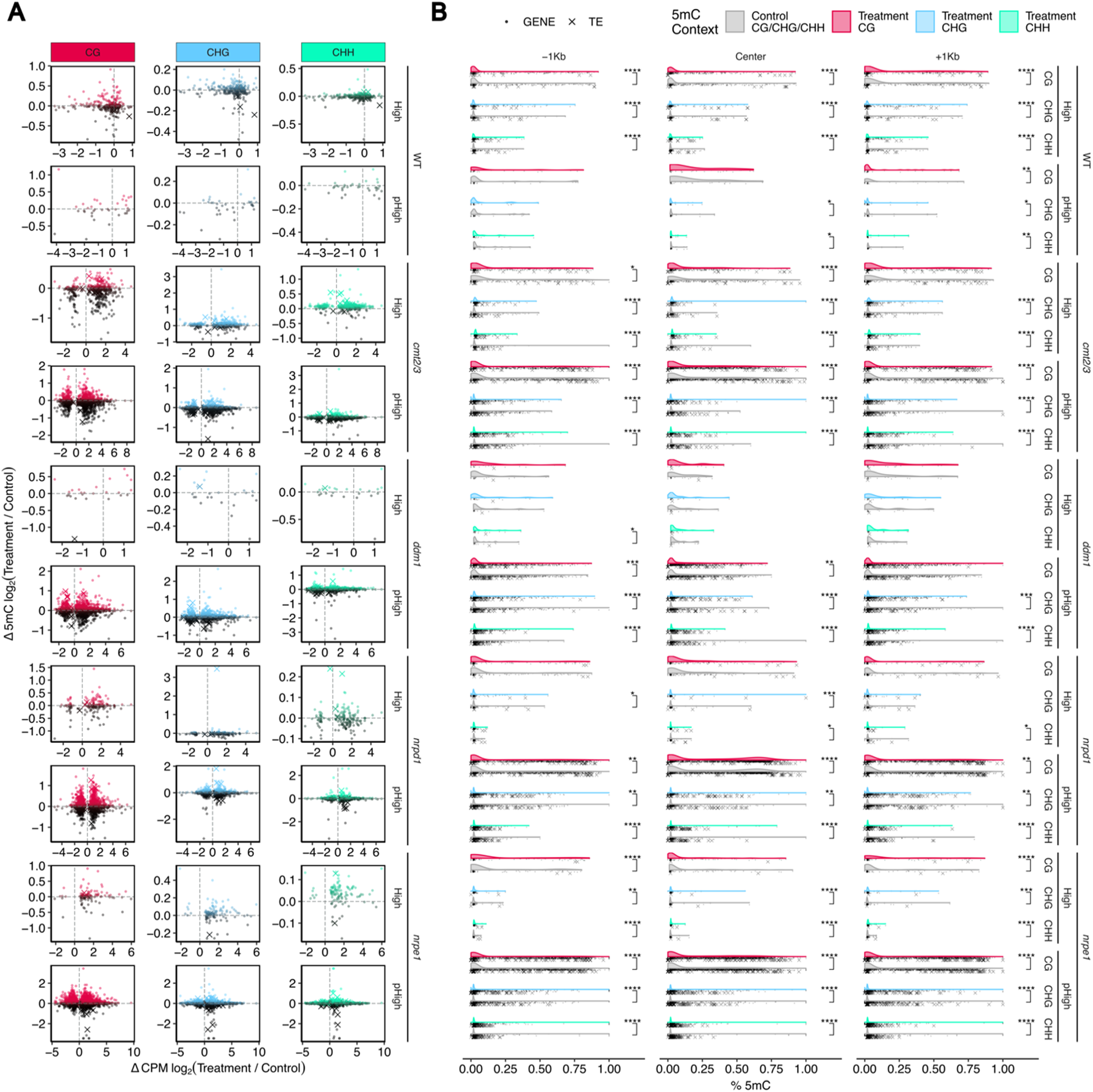
Differential transcript abundance of genes and TEs is associated with changes in DNA methylation across sequence contexts. (A) Scatter plots showing the relationship between changes in mean expression level (Δlog_2_(CPM), x-axis; Treatment/Control) and log2 fold-change in transcript abundance (Δlog_2_FC, y-axis) for differentially abundant genes (DAGs; circles) and TE fragments (DATEs; crosses) across CG, CHG, and CHH methylation contexts (columns) in all five genotypes and CO_2_ treatment conditions (rows; High and pHigh relative to paired Control). Points are colored by direction and significance of differential abundance (red = significantly upregulated; blue = significantly downregulated; grey = not significant; FDR < 0.05, *P* < 0.05, |log2FC| ≥ 1). The distribution of points along the x-axis reflects the association between baseline expression level and the magnitude of CO_2_-dependent transcriptional change, while the y-axis captures the directionality and extent of differential abundance within each methylation-sequence context. (B) Bee swarm distributions of fractional 5-methylcytosine (5mC) levels (% 5mC, x-axis; 0–1) at DAG and DATE loci across all three methylation sequence contexts (CG, CHG, CHH; columns) and CO_2_ conditions (Control, Treatment; colored by context as indicated in the legend), stratified by genotype (rows). Each point represents one locus; horizontal bars indicate the median. The distribution of 5mC values at DAG and DATE loci across conditions and genotypes is consistent with CO_2_-associated redistribution of DNA methylation at differentially regulated loci, with patterns varying by sequence context and genetic background. In WT plants, DATEs show increased transcript abundance coincident with loss of 5mC across all sequence contexts under elevated CO_2_ (Fig. 5D), a pattern that is absent or attenuated in CMT2/3-deficient backgrounds, consistent with a role for non-CG methylation maintenance in TE silencing. Methylation quantification was performed using EM-seq data as described in the Methods.

**Supplementary Figure S8.**
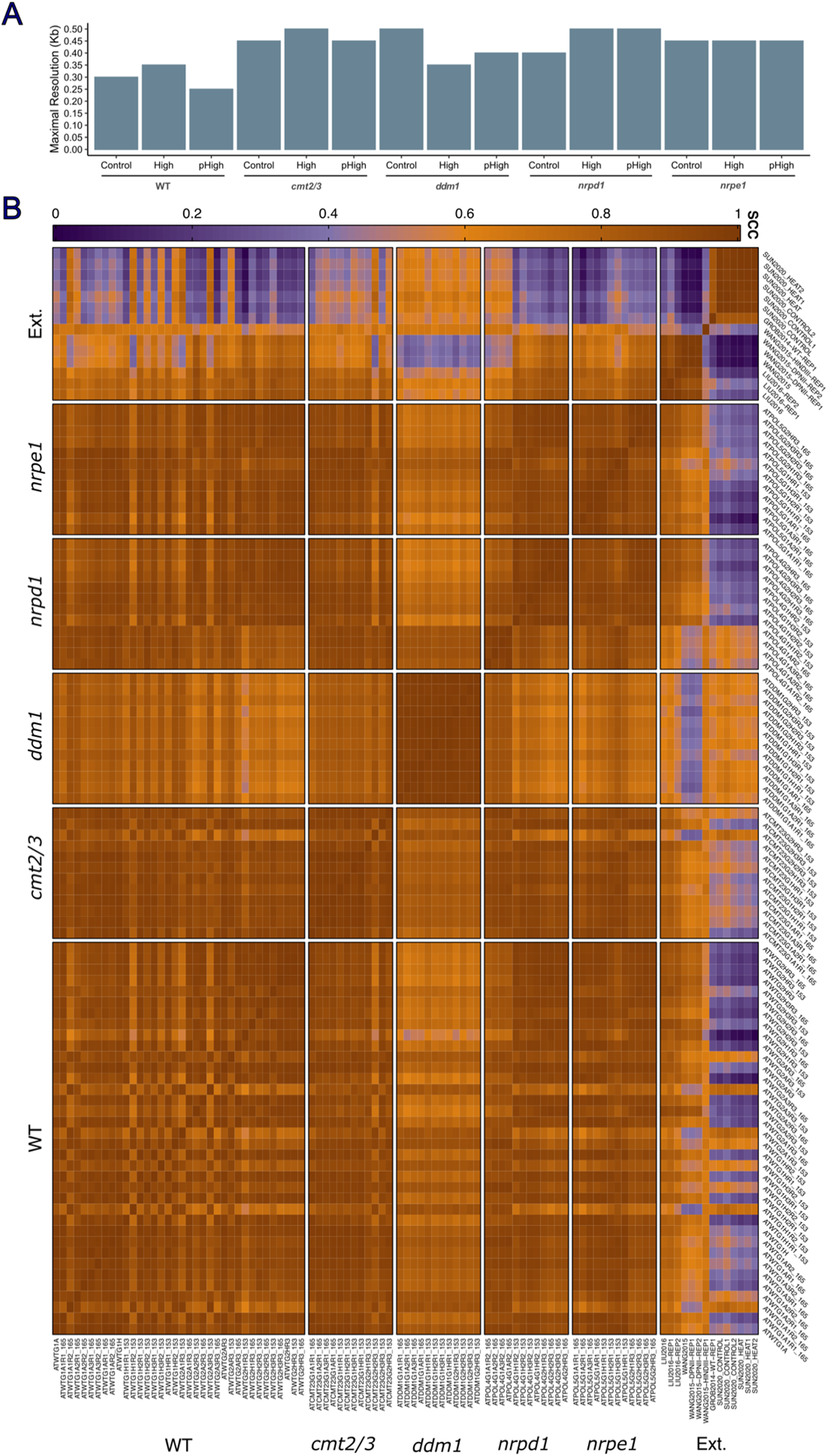
Hi-C libraries achieve gene-level resolution and high inter-library concordance. **(A)** Maximal resolution (Kb) achieved by merged Hi-C contact matrices for each genotype (WT, *cmt2/3*, *ddm1*, *nrpd1*, *nrpe1*) and CO_2_ treatment condition (Control, High, pHigh). Maximal resolution was determined as the finest bin size with ≥ 80% coverage and at which ≥1,000 contacts per bin were recovered genome-wide following KR normalization (Rao et al., 2014). All libraries exceeded 1 Kb maximal resolution after replicate merging. Bar heights reflect the maximal resolution obtained via replicate consensus of unique reads per condition. **(B)** Pairwise Stratum Adjusted Correlation Coefficient (SCC) matrix (Yang et al., 2017) calculated between all individual Hi-C replicate libraries from this study and published external (Ext.) *Arabidopsis* Hi-C libraries. Color scale indicates SCC value (orange/yellow = high concordance; purple = lower concordance). Within-genotype replicates show high pairwise SCC values, confirming biological reproducibility. SCC values between libraries of the same genotype and condition are consistently higher than those across conditions or between this study and external datasets, reflecting genuine biological differences in chromatin organization across CO_2_ treatments and genetic backgrounds rather than technical variation. The lower SCC values observed between internal and external libraries are consistent with differences in experimental protocol, tissue type, developmental stage, or sequencing depth between studies. External datasets used for comparison referenced throughout the main text.

**Supplementary Figure S9.**
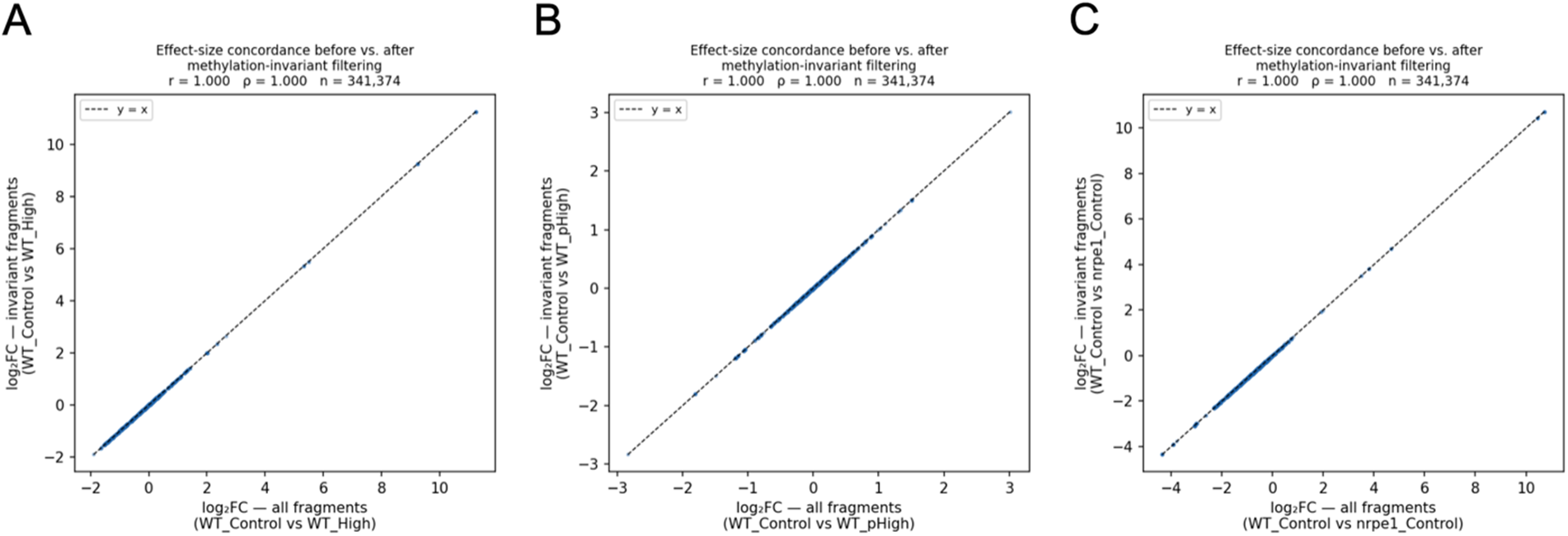
Methylation-invariant fragment analysis supports the robustness of Hi-C differential contact frequency estimates against Sau3AI methylation-dependent digestion bias. Effect-size concordance scatter plots comparing log2 fold-change in contact frequency (log2FC) computed from all Sau3AI fragments (x-axis) against log_2_FC computed from the subset of methylation-invariant fragments (y-axis) for three pairwise comparisons: (A) WT Control vs. WT High, (B) WT Control vs. WT pHigh, and (C) WT Control vs. *nrpe1* Control. Methylation-invariant fragments (n = 341,374; 78.4% of all 435,342 Sau3AI fragments, covering 78.1% of the genome) were defined as fragments whose boundary sites showed fractional methylation below 10% across all samples and conditions in EM-seq data, with a minimum 5x read coverage. Effect sizes were computed as described in the Methods. In all three comparisons, effect sizes are highly concordant between the full-fragment and invariant-fragment analyses (Pearson r >0.9999, Spearman ρ >0.9999, n = 341,374), and the log2FC distributions of invariant and excluded fragments were similar (mean log_2_FC 0.068 vs. 0.045; KS statistic D = 0.175, p < 0.001), indicating that fragments excluded due to differential methylation-dependent cut-site recovery do not show a systematically different CO_2_-associated contact response. Together these results support the interpretation that differential Sau3AI digestion is not a primary driver of the chromatin contact frequency differences reported in this study. The three comparisons shown represent the primary CO_2_ response (A), the transgenerational comparison (B), and the genotype with the most severely disrupted methylation landscape (C), constituting the comparisons most vulnerable to the potential confound that this analysis addresses.

**Supplementary Figure S10.**
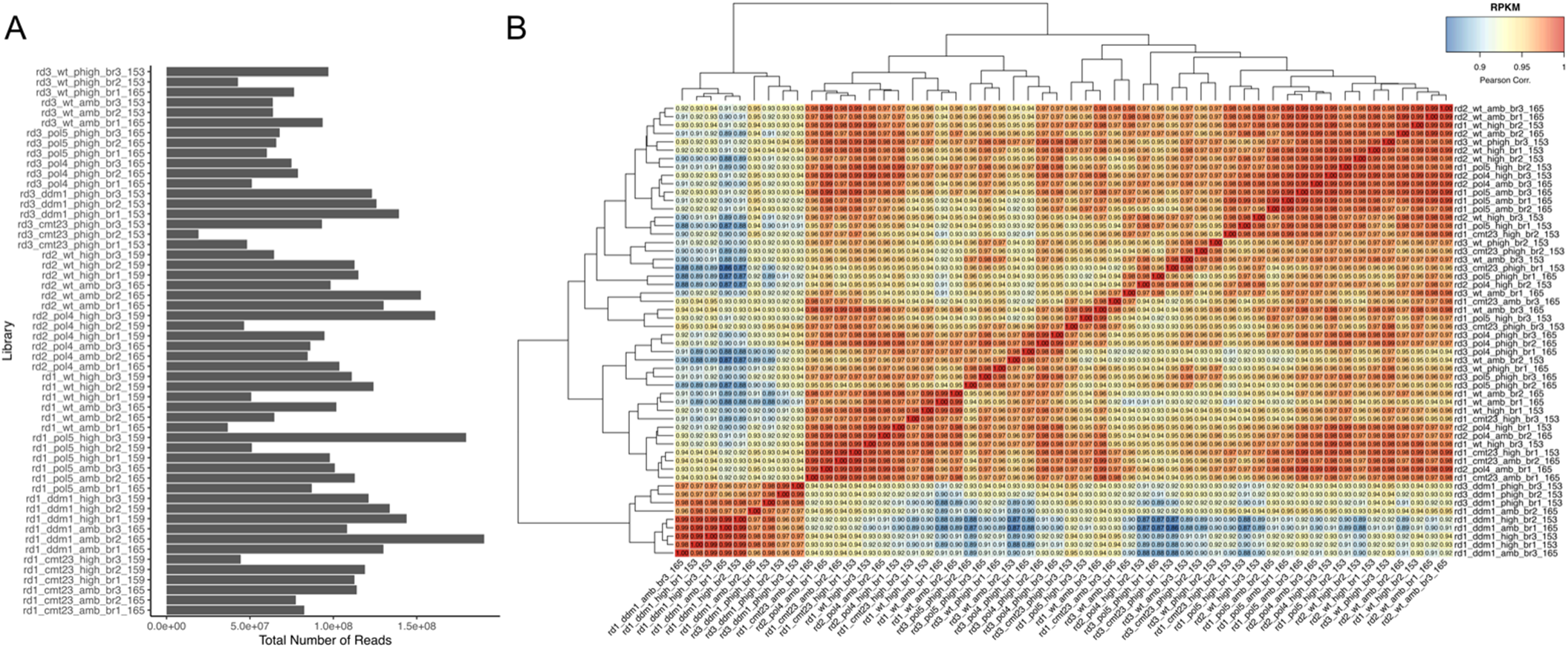
Quality control and reproducibility assessment of RNA-seq libraries. **(A)** Total number of mapped reads per RNA-seq library across all genotypes (*cmt2/3*, *ddm1*, *nrpd1*, *nrpe1*, and WT) and CO_2_ treatment conditions (Control, High, and pHigh). Libraries are ordered by sample identifier on the y-axis. All libraries exceeded the minimum depth threshold required for differential abundance analysis. **(B)** Pairwise Pearson correlation matrix of normalized transcript abundance (RPKM) across all RNA-seq libraries, with hierarchical clustering applied to both rows and columns. Color scale indicates the correlation coefficient strength (red = high; blue = low). Libraries cluster primarily by genetic background and secondarily by CO_2_ treatment condition, consistent with genotype as the dominant source of transcriptional variation. Within-condition and within-genotype replicates show high pairwise correlations, confirming library reproducibility. The clustering structure also reflects the known relationships between the RdDM pathway mutants (*nrpd1*, *nrpe1*) and maintenance methylation mutants (*cmt2/3*, *ddm1*), with WT libraries forming a distinct cluster corresponding to each treatment condition. Batch effects due to growth round and chamber were incorporated as blocking factors in the differential expression model (see Methods).

**Supplementary Figure S11.**
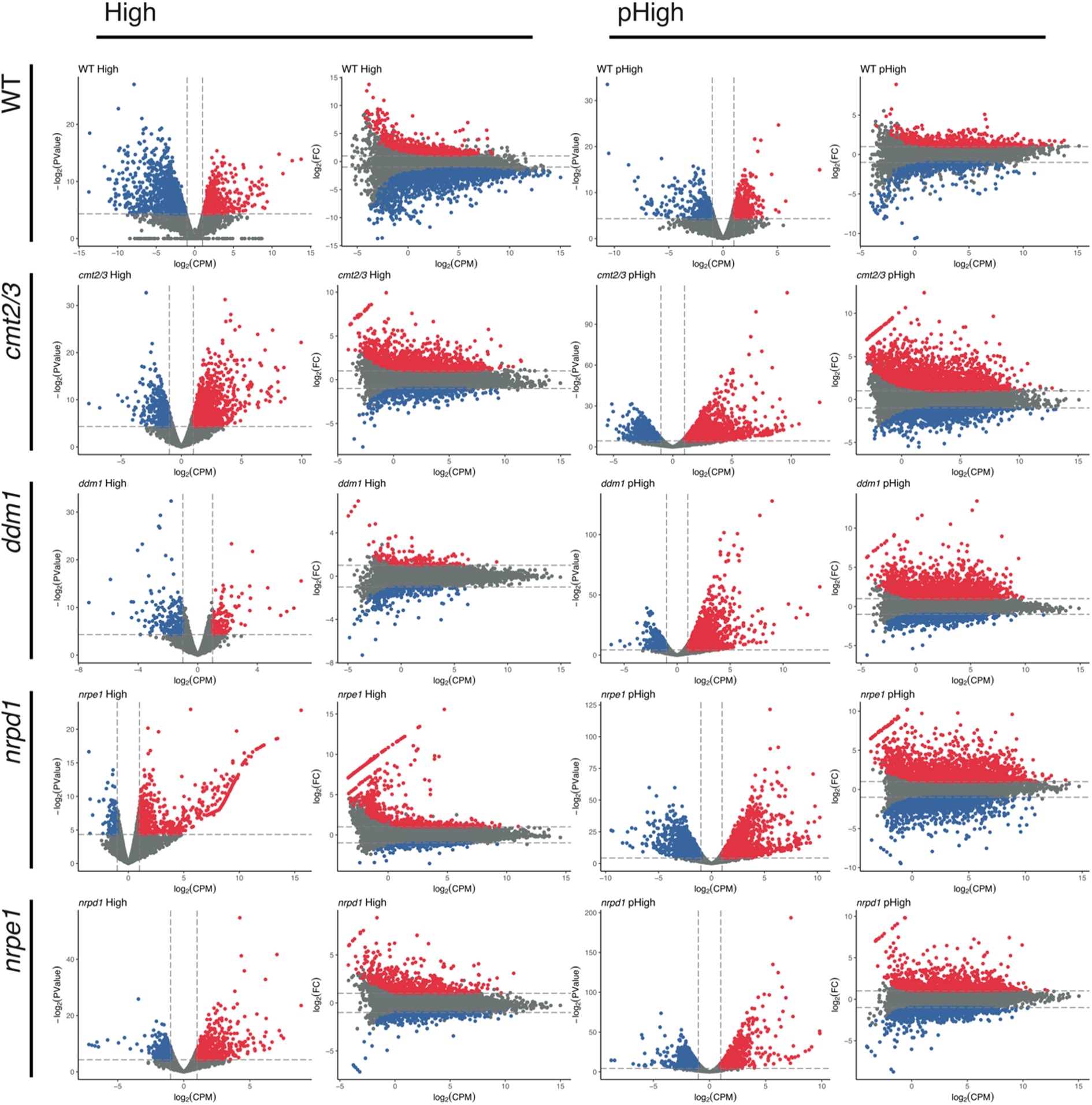
Genome-wide differential abundance of genes and transposable elements across CO_2_ treatments and genetic backgrounds. Volcano (left) and MA (right) plots for each genotype and generation showing GLM residual p-values (-log2(PValue), y-axis) or log_2_ fold-change (log2FC, y-axis) as a function of mean expression level (log(CPM), x-axis) for differentially abundant genes and TEs in High CO_2_ and progeny-of-High (pHigh) plants relative to paired ambient CO controls, across all five genetic backgrounds (rows) examined: wild type (WT), *cmt2/3*, *ddm1*, and *nrpe1*, *nrpd1*. Red points indicate significantly upregulated loci (log2FC ≥ 1, FDR < 0.05, *P* < 0.05); blue points indicate significantly downregulated loci (log2FC ≤ −1, FDR < 0.05, *P* < 0.05); grey points indicate loci that did not meet significance thresholds. Dashed horizontal lines mark the minimum log2FC ≥ |1| threshold. Differential abundance was assessed using a multifactorial generalized linear model incorporating growth round and chamber as blocking factors to control for batch effects, with normalization and statistical testing performed using EdgeR (Chen et al., 2025). The complete list of significant DAG and DATE accessions across all genotypes and conditions is provided in Supplementary Data S2.

